# A Bayesian Hierarchical Model for Signal Extraction from Protein Microarrays

**DOI:** 10.1101/2022.02.16.480698

**Authors:** Sophie Bérubé, Tamaki Kobayashi, Amy Wesolowski, Douglas E. Norris, Ingo Ruczinski, William J. Moss, Thomas A. Louis

**Affiliations:** Department of Biostatistics, Johns Hopkins University Bloomberg School of Public Health, 615 N. Wolfe Street Baltimore, MD, USA

**Keywords:** Bayesian models, Bioinformatics, Optimal ranks, Pre-Processing, Protein microarrays

## Abstract

Protein microarrays are a promising technology that measure protein levels in serum or plasma samples. Due to the high technical variability of these assays and high variation in protein levels across serum samples in any population, directly answering biological questions of interest using protein microarray measurements is challenging. Using within-array ranks of protein levels for analysis can mitigate the impact of between-sample variation on downstream analysis. Although ranks are sensitive to pre-processing steps, ranking methods that accommodate uncertainty provide robust and loss-function optimal ranks. Such ranking methods require Bayesian modeling that produces full posterior distributions for parameters of interest. Bayesian models that produce such outputs have been developed for other assays, for example DNA microarrays, but those modeling assumptions are not appropriate for protein microarrays. We develop and evaluate a Bayesian model to extract a full posterior distribution of normalized fluorescent signals and associated ranks for protein microarrays, and show that it fits well to data from two studies that use protein microrarrays from different manufacturing processes. We validate the model via simulation and demonstrate the downstream impact of using estimates from this model to obtain optimal ranks.

## 1. Introduction

Protein microarrays are assays that quantify levels of antibodies and other proteins in human or animal serum or plasma. They are versatile and have been employed in numerous areas of research such as cancer therapeutics, infectious disease diagnostics and biomarker identification (Huang and Zhu, 2017; Zhu *and others*, 2006; Nagele *and others*, 2011; Ramachandran *and others*, 2008; Hartmann *and others*, 2009). However, measurements in protein microarray studies contain high levels of between-sample variation, that may obfuscate biological signals of interest even after preprocessing steps like normalization (Brezina *and others*, 2015; Sboner *and others*, 2009; Bérubé *and others*, 2021). Ranking proteins according to their levels of fluorescent intensity measures the quantity of protein in a sample relative to quantities of other proteins in that sample thereby minimizing the impact of between-sample variability and enabling analysts to directly answer scientific questions of interest. Biological inferences can then be made across samples using these within-sample ranks.

Accommodating uncertainty in estimating ranks by optimizing a loss function produces values that out-perform competing approaches and Bayesian formalism is particularly effective in this regard (Shen and Louis, 1998; Lin *and others*, 2006; Henderson and Newton, 2016). In the micro-array context, ranks report relative quantities of target molecules across samples, a relevant biological signal (Gu *and others*, 2014; Cantarel *and others*, 2014; Shiraishi *and others*, 2013). Empirical Bayes models have been used in DNA microarray analyses to improve the ranks of normalized gene expression levels, and maximize the sensitivity in selecting differentially expressed genes with the largest effects (Noma *and others*, 2010; Efron *and others*, 2001; Gottardo *and others*, 2003). However these models rely heavily on the assumption that the majority of genes investigated have similar expression levels across samples. Protein levels are highly variable across individuals for reasons are not fully understood; therefore it is unreasonable to assume that the majority of proteins investigated in a protein microarray study are present at similar levels across samples.

We develop and evaluate a Bayesian model that extracts full posterior distributions for normalized fluorescent intensity from protein microarrays, and the availability of these distributions enables improved rankings and other inferences. Our model uses information from repeated measurements and controls across the assay to inform an error structure that is then subtracted from observed measurements to obtain an estimate of true underlying fluorescent signal, resulting from specific protein binding. We show that the model, with some simplifications to improve computational efficiency, fits well to two previously published protein microarray studies: (Kobayashi *and others*, 2019) and (Pan *and others*, 2017). Each study uses protein microarrays with different numbers of probes, sets of proteins, and manufacturing procedures, suggesting that our model has the potential for applications to a wide variety of protein microarray studies. We show the impact of our model on downstream analysis by comparing protein ranks obtained from different ranking and pre-processing steps. Finally, we validate results with a simulation study. This model provides full posterior distributions for estimates of the quantity of target protein in a sample and enables the use of ranking procedures that incorporate uncertainty. These methods yield optimal protein ranks that support a variety of downstream analyses and enable robust and accurate biological inference.

## 2. Notation and Methods

### 2.1 Protein Microarray Components

Protein microarrays are comprised of proteins spotted on glass slides and organized into rows and columns. A serum or plasma sample is added to the slide and antibodies or other proteins of interest in this sample bind to specific proteins on the array. Using fluorescent tags, binding events on the array are detected by a scanner that produces two observable signals. These are background signal, which is a measure of the fluorescent intensity in an outer ring surrounding a particular spot, and foreground signal which is the measure of the fluorescent intensity at the center of a particular spot (Zhu and Snyder, 2001). Protein array-based studies seek to characterize proteins in multiple human or animal samples, meaning that the number of samples analyzed is typically equal to the number of arrays in the study. We denote fluorescent intensity measurements from the scanner in a particular study with the notation *R*_*i,t*(*p*),*p*(*i,j*),*j*_ or its corresponding transformed measurement *Y*_*i,t*(*p*),*p*(*i,j*),*j*_, with the following subscripts:

- array number *i ∈* {1, …, *I*}
- spot *j ∈* {1, …, *J*_*i*_}, where *J*_*i*_ is the number of spots on array *i* in a study, since the number of spots on an array does not usually change within a study, we assume that *J*_*i*_ is the same for all *i ∈* {1, …, *I*}.
- protein *p*(*i, j*) *∈* {1, …, *P* }, each individual spot *j* is spotted with a protein (or buffer if it is a negative control).
- spot type *t*(*p*), we consider three broad categories of proteins, negative controls, positive controls, and active spots, and within these three categories there are multiple types of spots, *t*(*p*), that depend on the protein (or lack of protein in the case of negative controls). For the control protein types (negative and positive controls), *t*(*p*) *∈* {1, …, *T* } and for active spots *t*(*p*) *∈* {*a*_1_, …, *a*_*A*_}. The designations of *t*(*p*) for the HuProt^*T M*^ and malaria arrays are shown in Tables S1 and S2 respectively.

The two protein microarrays used for each study have different compositions. The HuProt^*T M*^ arrays used by Pan *and others* (2017) contain 43776 active probes corresponding to approximately 16000 different proteins (each with multiple replicates on an array) chosen to provide broad coverage of the known human proteome. We consider all these active proteins to be of the same type, *t*(*p*) = *a*_1_. The arrays also contain 4608 control proteins corresponding to multiple replicates of 17 types (*T* = 17) of negative and positive controls. In their study, Pan *and others* (2017) analyze the antibody profiles of 100 individuals. Of these individuals, 20 were healthy and the remaining 80 had a form of lung cancer.

The arrays used by Kobayashi *and others* (2019) contain 500 *Plasmodium falciparum* and 500 *P. vivax* specific antigens referred to as active proteins of the same type, *t*(*p*) = *a*_1_ as well as 17 types (*T* = 17) of positive and negative control proteins. In this study, 429 samples from 290 individuals were collected across three sites in malaria endemic regions of Zambia and Zimbabwe. Across all three study sites, a random stratified sampling scheme was employed for household selection and every individual present in the household at the time of visit was eligible for enrollment meaning that not all individuals in the study had malaria at the time of sampling. For each individual enrolled in the study, at least one serum sample was collected and spotted on an array. Additionally, some arrays were spotted with sera from adults residing in the USA who had never traveled to a malaria endemic region to serve as controls.

### 2.2 Normalized Data

We use the output of the pre-processing pipeline described in Bérubé *and others* (2021) as the data that will be fed into the Bayesian model. Briefly, the ratio of observed foreground signal to background signal at each probe (*Y* ^*′*^) is log transformed. Then, using a robust linear model described by Sboner *and others* (2009), array and subarray effects are estimated and subtracted from log(*Y* ^*′*^) to produce 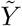. Finally, these values are standardized using array specific sample means and sample standard deviations of control probes to produce *Y*, the input for the Bayesian Model. These starting values for the malaria data are available on a public Github repository (https://github.com/sberube3/Protein_array_bayesian_model).

### 2.3 Bayesian Models

#### 2.3.1 Complete Model

The following model describes the relationship among observed and then transformed data, *Y* as described in Bérubé *and others* (2021) and true underlying signal *S*. We assume that true underlying fluorescent intensity resulting from specific protein binding between the proteins in the sample and the probes on the array, *S*, adds to error *e*, which is the technical variation remaining in the measurements after the pre-processing pipeline (Bérubé *and others*, 2021), to produce the normalized observation *Y*. Equation 2.1 is the full hierarchical Bayesian model describing this relationship. In order to leverage the control proteins present on all arrays to inform the distribution of the errors *e*_*i,t*(*p*),*p*(*i,j*),*j*_ for *t*(*p*) *∈* {1, …, *T* }, we consider the empirical distributions of mean centered control probes 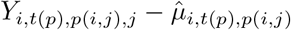 where 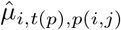 is the sample mean taken over all spots of type *t*(*p*), *∈* {1, …, *T* } on each array. The observed densities of *e*_*i,t*(*p*),*p*(*i,j*)_ for *t*(*p*) *∈* {1, …, *T* } (Figure S1) are similar across all proteins, therefore it is reasonable to assume that the errors in Equation 2.1 are identically distributed, and well approximated by a generalized beta-generated distribution (Alexander *and others*, 2012).

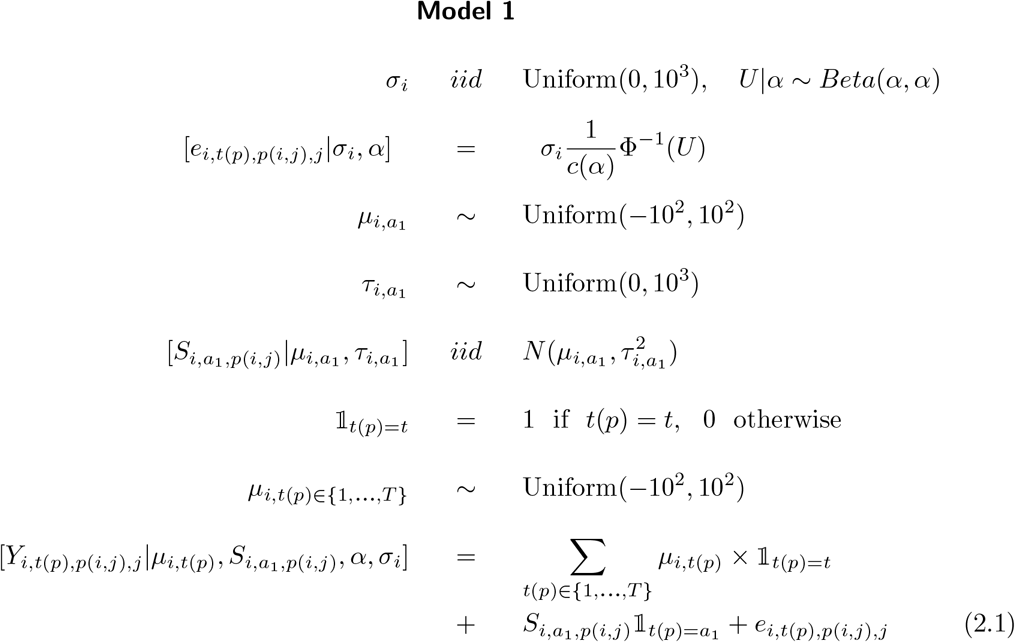

Importantly, *c*(*α*) is a constant, used to ensure that the standard deviation of *e*_*i,t*(*p*),*p*(*i,j*),*j*_ is approximately *σ*_*i*_ for an array *i*, and we approximate *c*(*α*) in the following way:

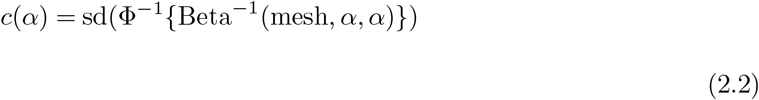

where the mesh is a sequence of values between 0 and 1 increasing by increments of 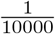. We also use a numerical approximation procedure to estimate the maximum likelihood value of *α* given *Y*_*i,t*(*p*),*p*(*i,j*),*j*_ for *t*(*p*) *∈* {1, …, *T* } at 2.5, Appendix B details this procedure.

#### 2.3.2 Simplified Models

Additionally, we propose and evaluate two simplifications to the complete model in Equation 2.1 to increase computational efficiency. First, letting *α* = 1 we obtain Model 2, which is equivalent to assuming a Gaussian distribution for the errors.

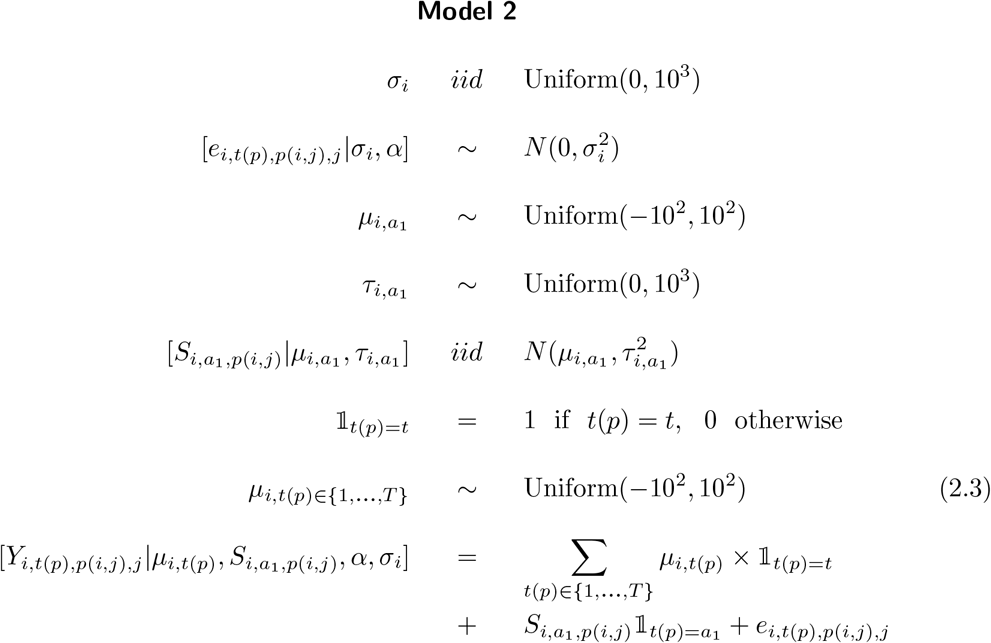

Second to obtain Model 3 we use an empirical point estimate of *σ*_*i*_, the variance of the error distribution, 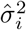:

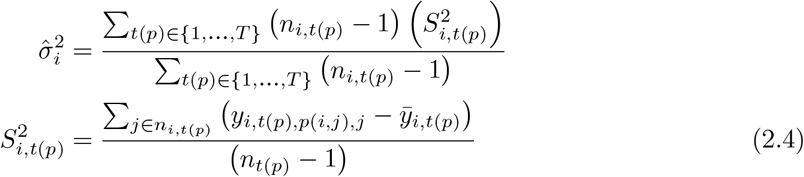

where *n*_*i,t*(*p*)_ is the number of spots of type *t*(*p*) on an array *i* and 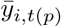 is the mean of *y*_*i,t*(*p*),*p*(*i,j*),*j*_ across all spots of type *t*(*p*) in array *i*.

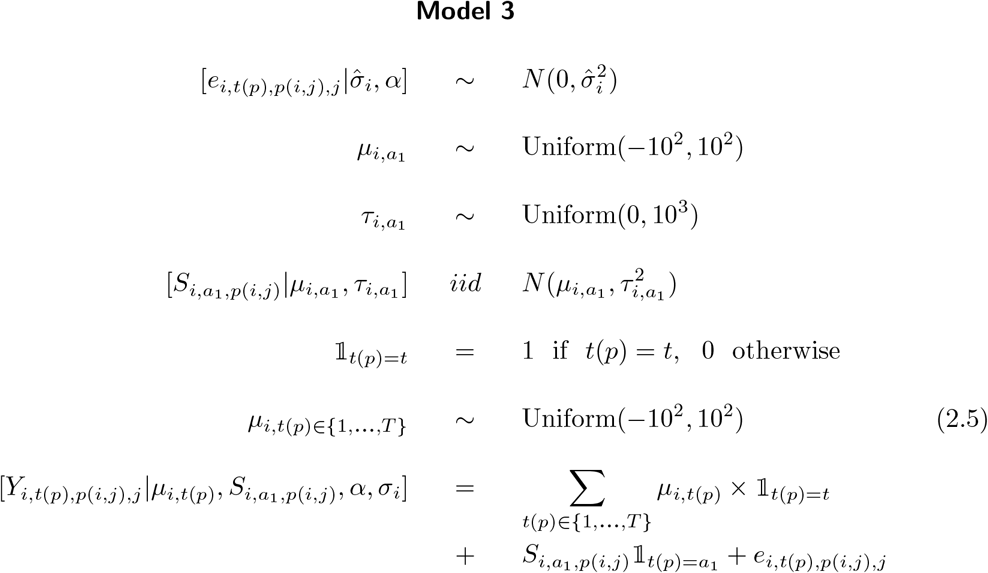

#### 2.3.3 Estimation Procedure

We aim primarily to estimate the full posterior distribution of normalized fluorescent intensity, *S*_*i,t*(*p*),*p*(*i,j*)_|*Y*_*i,t*(*p*),*p*(*i,j*)_ for all active spots on each array, and also evaluate the full posterior distributions of the other parameters, namely, *μ*_*i,t*(*p*)_ for *t*(*p*) ∈ {1, …, *T*}, 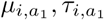, and *σ*, where relevant. We use Markov Chain Monte Carlo (MCMC) estimation implemented through the Rjags package. We run each array *i* in a study independently but jointly estimate all parameters for all spots *j* and proteins *p*(*i, j*) within each array. For each array, we use a burn-in of 5,000 draws, and posterior distributions are estimated using 10,000 draws after burn-in with no thinning.

#### 2.3.4 Evaluating Model Fit

We evaluate the fit of our model using percentile based residuals mapped to the associated quantile of a standard Gaussian distribution (Bérubé *and others*, 2019; Yan and Sedransk, 2007). Specifically, for each *y*_*i,t*(*p*),*p*(*i,j*),*j*_ in our dataset, we compute the transformed residual 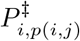, as follows:

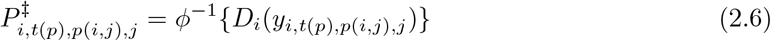

where *D*_*i*_ is the combination of posterior distributions across all *S*_*i,p*(*i,j*)_ and *D*_*i*_(*y*_*i,t*(*p*),*p*(*i,j*),*j*_) is the percentile location of the observed *y*_*i,t*(*p*),*p*(*i,j*)*j*_ in *D*_*i*_.

### 2.4 Simulation Setup

We conducted a simulation study to validate Models 1, 2, and 3 with three steps: data generating, data analysis, and evaluating model fit and ranking procedures. For the data generating step, we generate data from three different models. Data A and D in which data are generated from Model 1 (Equation 2.1) where *σ*_*i*_ is drawn from a Uniform(0, 10) distribution instead of a Uniform(0, 10^3^) distribution, in Data A the number of active and control probes roughly matches those on the malaria arrays and in Data D the number of active and control probes roughly matches those on the HuProt^*T M*^ arrays. Data B and E, which are generated from Model 2 (Equation 2.3), also with *σ*_*i*_ is drawn from a Uniform(0, 10) distribution, in Data B the number of active and control probes roughly matches those on the malaria arrays, and in Data E, the number of active and control probes roughly matches those on the HuProt^*T M*^. Data C and F, which is generated from Model 2 (Equation 2.3), with *σ*_*i*_ is drawn from a Uniform(0, 10^3^) distribution, in Data C the number of active and control probes roughly matches those on the malaria arrays, and in Data F the number of active and control probes roughly matches those on the HuProt^*T M*^ arrays. The full code for this can be found on Github (details in Section 5). For the data analysis step, we fit Models 1 and 2, exactly as they appear in Section 2.3 to data sets A, B, C, D, E, and F. Finally, we assess model fit under these data generating and analysis scenarios by using the percentile residuals *P*^‡^ (Section 3) and compare the ranks computed using the posterior distributions *S*|*Y* to the true ranks of *S*.

### 2.5 Ranking Methods

Here we present two methods that use the full posterior distributions of signal (the output of our Bayesian model) to rank proteins on a microarray by their fluorescent intensities, both are described in Lin *and others* (2006).

#### 2.5.1 Method A, Squared Error Loss (SEL)

We define the ranks of the true *S*_*i,p*(*i,j*)_ to be:

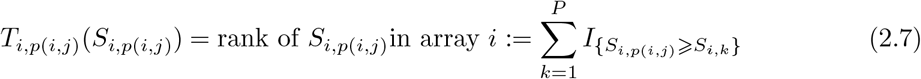

so that the smallest *S* will have rank 1 and the largest will have rank *P*. The estimator that minimizes squared error loss is the posterior mean:

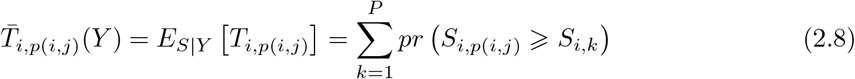

We use burned in MCMC draws to compute this quantity by computing the rank of each protein in each array for each MCMC draw, and then averaging those ranks across the MCMC draws. We compute 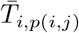 as well as the integer version of these:

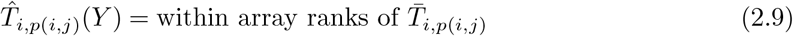

We report these values, combined across arrays, specifically the simple average of 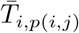 across arrays:

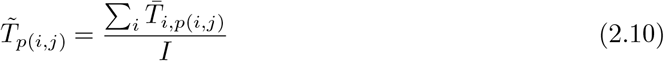

As well as the integer version of these across arrays:

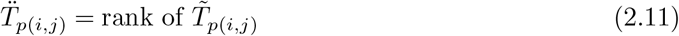

#### 2.5.2 Method B, Classification with above γ Loss

We consider an estimator that will allow us to identify the top *γ*% of proteins by fluorescent intensity within arrays. Specifically we let:

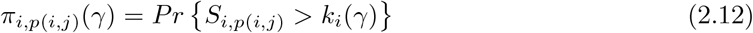

where MCMC draws are indexed by *ν ∈* {1, …, *M* } and where *k*_*i*_(*γ*) is the value in the empirical distribution of *S*_*i,p*(*i,j*),*ν*_ over all *p*(*i, j*), *ν* in array *i* that corresponds to the (100 *− γ*) percentile. We estimate the value 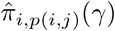 by defining the following indicator function:

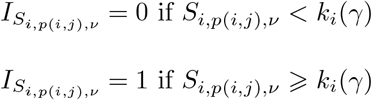

and therefore:

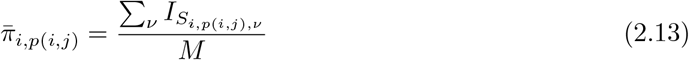

We also consider the ranks of these *π* values:

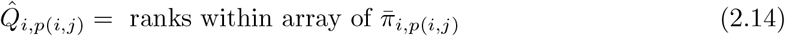

We also define:

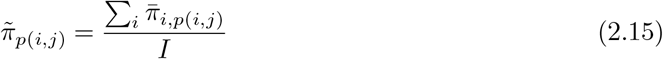

And the integer ranks of these values:

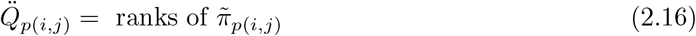

As an example, we show the case where *γ* = 3 in our analysis. In the malaria arrays the top 3% of proteins corresponds to roughly to the top 30 proteins by fluorescent intensity, in the HuProt^*T M*^ arrays this corresponds roughly to the top 450 proteins by fluorescent intensity.

All code for the methods outlined in Sections 2.3.3, 2.3.4, 2.4, 2.5.1,and 2.5.2 can be found on a public Github repository (https://github.com/sberube3/Protein_array_bayesian_model).

## 3. Results

We fit Model 1 (Equation 2.1) to the protein microarray data from the lung cancer (Pan *and others*, 2017) and malaria (Kobayashi *and others*, 2019) studies. We show that along with producing a good overall fit, posterior distributions of the parameter *S*_*i,t*(*p*),*p*(*i,j*)_ for active (*t*(*p*) = *a*_1_) proteins, which estimate the quantity of target proteins in the sample, reveal clear separation across proteins with different corresponding *Y* values while also displaying shrinkage towards the mean 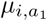, making them good candidates for downstream inference. We show that the simplifications in Models 2 and 3 (Equations 2.3 and 2.5), reduce computation time, provide similar posterior distributions for *S*_*i,t*(*p*),*p*(*i,j*)_ to those obtained with Model 1, and fit well to both datasets. Finally, we validate our results with a simulation study. The R code to execute all of these steps is available in a GitHub repository (details in Section 5).

### 3.1 Estimation and Fit Using Model 1

The posterior distributions for the estimates of true fluorescent signal in the active spots, *S*_*i,t*(*p*)=*a*_1_,*p*(*i,j*),*j*_ obtained by fitting Model 1 to the HuProt^*T M*^ and malaria arrays, show good separation for proteins with different observed (*Y*) values in both datasets (Figure 1, these four arrays from each study are representative of all arrays). Additionally, the posterior distributions for *S* also reveal shrinkage towards the mean 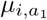 that is particularly pronounced for extreme values of *Y*, which is evidence that the model stabilizes these estimates, and provides good trade off between bias and variance for estimates of *S*. The wide uniform prior on *μ*_*i,t*(*p*)_ heavily weights the observations (*Y*) of repeated control probes in the estimation of the posterior mean *μ*_*i,t*(*p*)_|*Y* which allows the model to leverage information from repeated measurements on the array. Consequently, the posterior densities for the means of control proteins (*t*(*p*) *∈* {1, …, *T* }), *μ*_*i,t*(*p*)_ are centered exactly at the observed and transformed means *Y*_*i,t*(*p*),*p*(*i,j*),*j*_ for each type of control protein *t*(*p*) in both studies (Figure 2). Hyperparameters 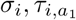, and 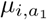 all show evidence of updating and trace plots for all parameters show good convergence (Figures S3, S4, S5, and S6). The observed quantiles of percentile-based residuals (*P*^‡^, Equation 2.6) for all arrays are close to those of a standard Gaussian distribution but with a few deviations showing the data has larger spread than what the models predict (Figure 3). This difference is fairly small though, suggesting a good overall fit for both studies.

**Fig. 1:**
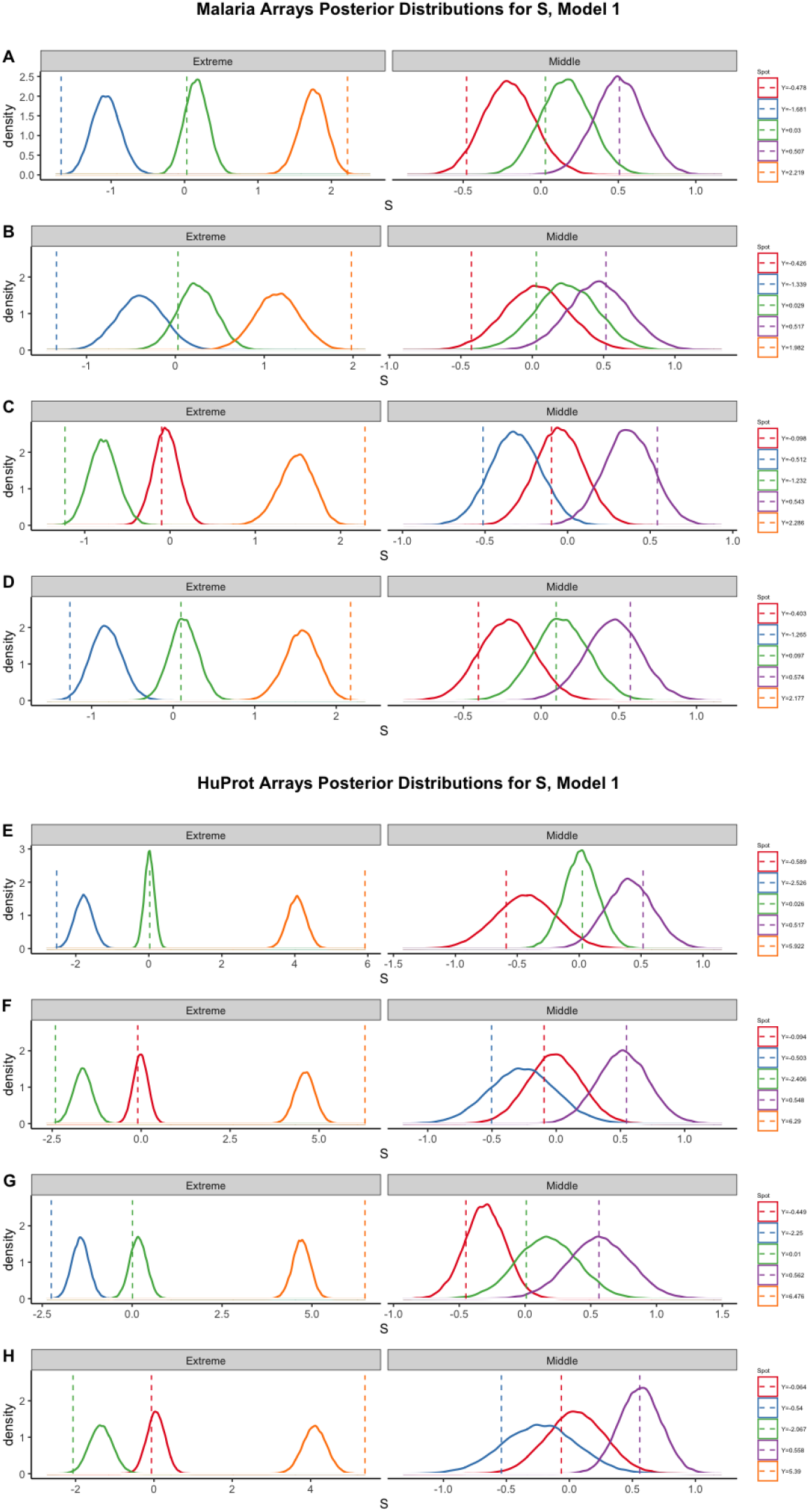
Posterior distributions for four different active spots (*t*(*p*) = *a*_1_) of *S* on four malaria arrays (A,B,C,D) and four HuProt^*T M*^ arrays (E,F,G,H) fit with Model 1. Vertical dashed lines are at mean of values *Y*_*i,t*(*p*)_=*a*_1,*p*(*i,j*),*j*_ corresponding to the protein *p*(*i, j*).

**Fig. 2:**
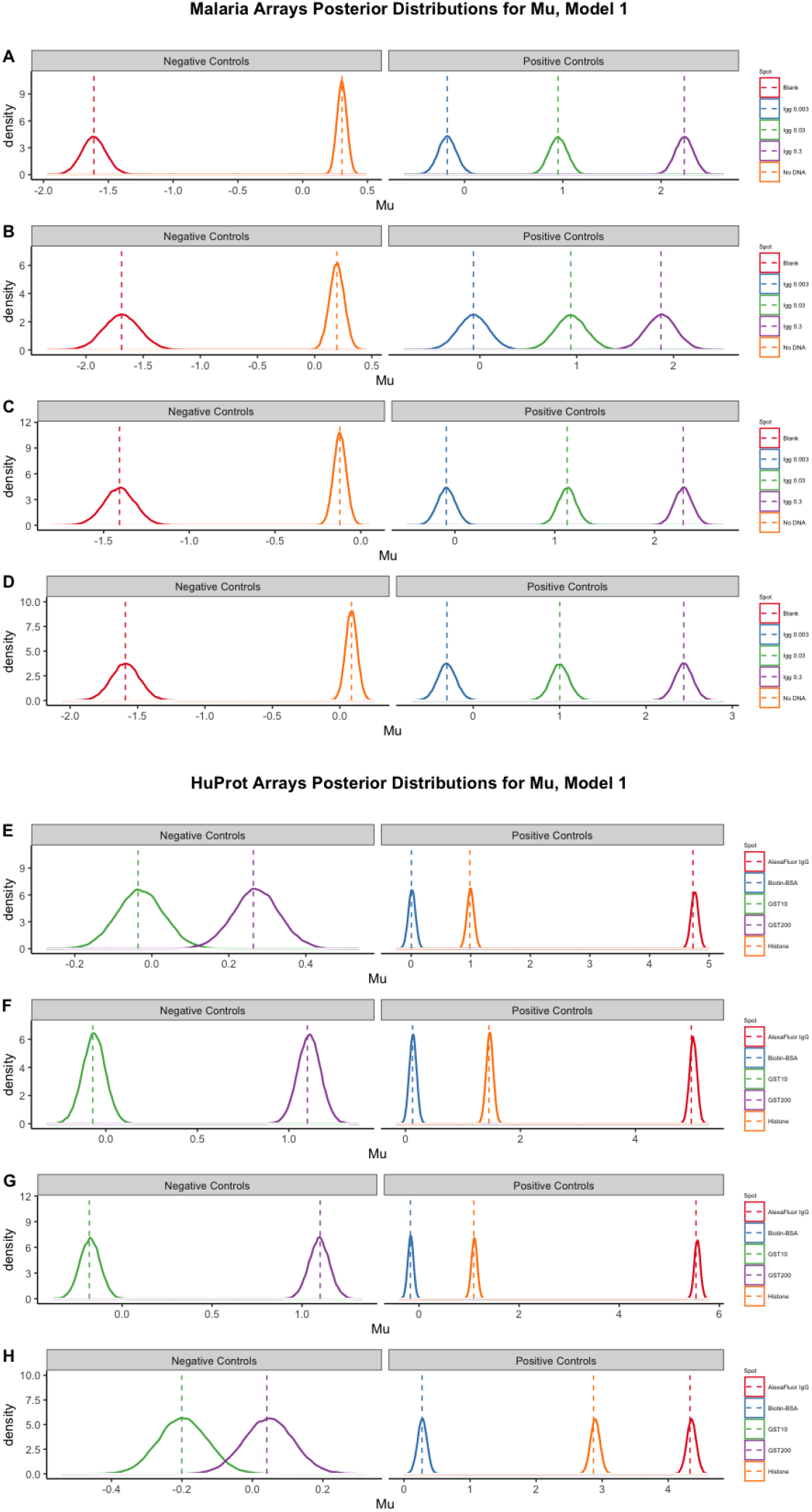
Posterior distributions for four different control spots *t*(*p*) *∈* {1, …, *T*} of *μ*_*i,t*(*p*)_ on four malaria arrays (A,B,C,D) and four HuProt^*T M*^ arrays (E,F,G,H) fit with Model 1. Vertical dashed lines are at mean of values *Y*_*i,t*(*p*),*p*(*i,j*),*j*_ corresponding to the control protein *p*(*i, j*).

**Fig. 3:**
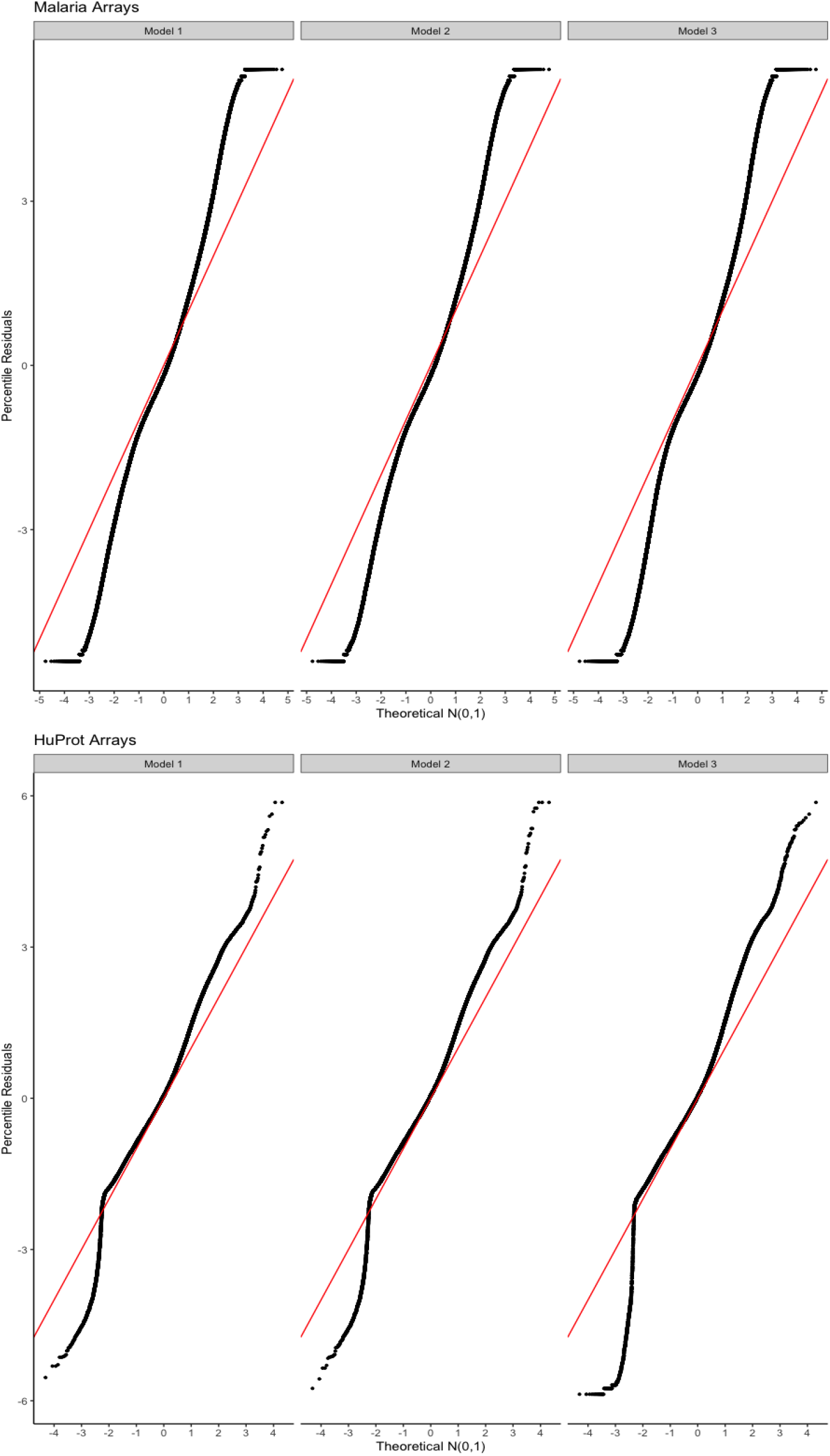
*P*^‡^ for all 503 malaria arrays and 5 of the 100 HuProt^*T M*^ arrays across Models 1, 2, and 3. The red line is *y* = *x*.

### 3.2 Estimation and Fit Using Models 2 and 3

Models 2 and 3 substantially reduce the computation time involved in obtaining full posterior distributions for all parameters, with Model 3 being slightly more efficient than Model 2 (Table 1), but both models show similar fit to both data sets when compared to Model 1. Despite the numerical approximation for *α* (Appendix B) suggesting that the empirical error distribution for signal on both protein microarray datasets (*e*_*i,t*(*p*),*p*(*i,j*),*j*_) is well approximated by a generalized Beta-generated distribution with *α* = 2.5, assuming a Gaussian distribution for the error structure (Models 2 and 3), produces similar overall fit to Model 1 in both data sets. Generally, the quantiles of percentile residuals, *P*^‡^ (Section 2.3.4) computed for *S* across all arrays in the malaria study and a subset of arrays in the lung cancer study match fairly closely to those of a standard Gaussian distribution across all three models with the data having greater spread than what the model estimates. Specifically, in the HuProt^*T M*^ array, the difference in spread between the data and model predictions are similar across Models 1 and 2 (Figure 3); Model 3 shows a larger difference, which could be attributed to the more extreme shrinkage of *S* posteriors towards the mean 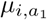 in Model 3 (Figures S7 and S8). In the malaria arrays, the shrinkage of *S* posteriors towards the mean 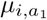 is fairly consistent across all three models, therefore all three models have similar differences between the spread of data and model predictions(Figure 3). Therefore the most computationally efficient model, Model 3, can be used instead of Model 1 with almost identical results for the malaria arrays, and Model 2, can be used with almost identical results in the HuProt^*T M*^ arrays. However, the differences in model fit between Models 1 and 3 in the HuProt^*T M*^ arrays are small and the posterior estimates of *S* from Models 2 and 3 achieve important goals for downstream analysis, such as good separation for proteins with different observed *Y* values, therefore it may still be worthwhile to consider fitting Model 3 to the HuProt^*T M*^ arrays with appropriate sensitivity analyses. Importantly, the posterior distributions for *μ*_*t*(*p*)_ in Models 2 and 3 look nearly identical to those of Model 1 for both the HuProt^*T M*^ and malaria arrays (Figures S9 and S10), hyperparameters show good updating and trace plots reveal good convergence (Figures S11-S18).

**Table 1:** Running times in minutes for the three models on individual malaria and HuProt^*T M*^ arrays (x1) as well as both datasets (x503 and x100 respectively).

| Model | Malaria Array (x1) | Malaria Data(x503) | HuProt<sup>TM</sup> (x1) | HuProt<sup>TM</sup> Data (x100) |
| --- | --- | --- | --- | --- |
| 1 | 9 | 4577 | 236 | 23600 |
| 2 | 2 | 956 | 52 | 5160 |
| 3 | 1 | 604 | 49 | 4880 |

### 3.3 Simulations

Percentile residuals for these simulations reveal good fit for generated data sets A, B, D, and E across analysis Models 1 and 2 (Figure 4) and ranks using posterior distributions of *S*|*Y* closely match generated values of *S* suggesting that the models are robust to the misspecification of the error distribution, providing further evidence in favor of using Models 2 and 3 which are more computationally efficient than Model 1. All eight scenarios involving data sets A and B show more extreme values having slight deviations from the expected standard Gaussian quantiles, this being most pronounced with data sets A and D (Figure 4) which may be due to the use of the Generalized-Beta distribution to estimate the error distribution. For generated data A and B the ranks obtained from posteriors using both analysis models 1 and 2 (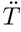 Section 2.5.1) match the true ranks of the top 30 values of *S* closely, and in both cases analysis models 1 and 2 produce identical ranks with the exception of one inversion in data A. The ranks obtained from posteriors using both analysis models 1 and 2 (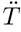 Section 2.5.1) fit to data C do not match the true ranks of *S* under this generating model (Figure 5). While the percentile residuals (*P*^‡^) for generated data C and F, fit with both Models 1 and 2, appear Gaussian (as evidenced by the fairly linear relationship between observed an expected quantiles), the deviations from the line *y* = *x* reveal that the variation in these residuals is higher than expected. This is due to simulated arrays with large values of *σ*_*i*_ (greater than 500), in these cases the model cannot accommodate the extreme variance in the observed data (*Y*), producing percentile residuals that deviate from the expected quantiles. Using information from the empirical distribution of control probes in both the malaria and lung cancer studies, such large values of *σ*_*i*_ seem unreasonable, especially after appropriate pre-processing and outlier identification. These simulations suggest that for most pre-processed data, the models are robust to error misspecification, the use of models the more computationally efficient models (2 and 3) is supported.

**Fig. 4:**
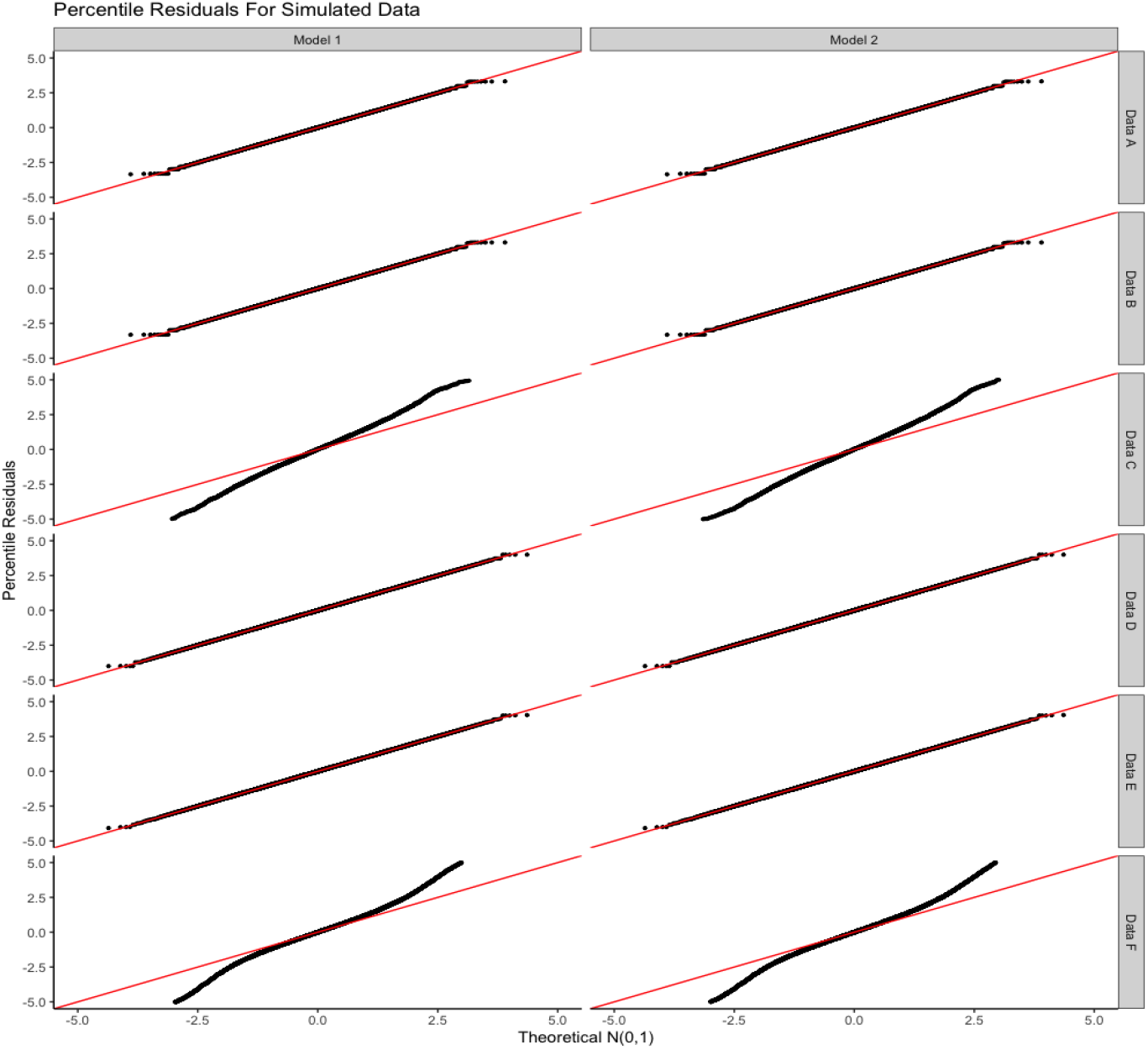
Percentile residuals (*P*^‡^) for 10 simulated arrays under generating models A, B, C, D, E, and F (Section 2.4), analyzed with Models 1 and 2 respectively.

**Fig. 5:**
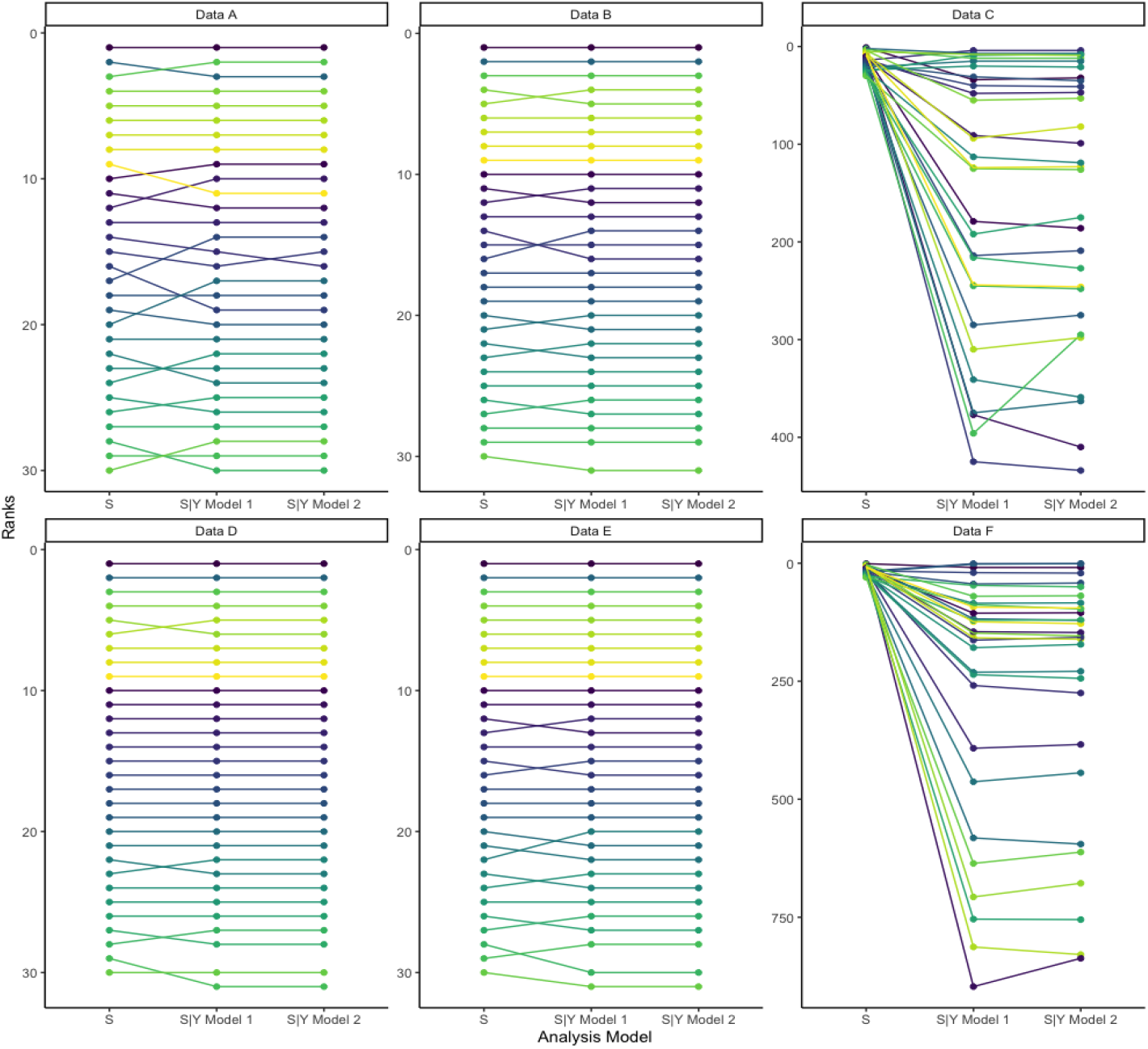
Each color is a protein simulated under generating mechanism *A, B, C, D, E*, or *F* each column shows one of the true ranks of *S*, the Squared error loss ranks 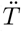, obtained from posterior distributions under Model 1 or Model 2. Lines represent how the protein ranks change from the top 30 true values to those for the two analysis models.

### 3.4 Impacts of Bayesian Modeling on Downstream Inference

Using the full posterior distribution of normalized fluorescent intensity, with two different loss functions (squared error loss, Section 2.5.1 and targeting top ranking proteins (the *γ*-ranks), Section 2.5.2) in each study produces different protein ranks (based on fluorescent intensity) from the previously published values. This result highlights the impact of using full posterior distributions of signals to produce optimal ranks on downstream inference. For example, Figure 6 shows how ranks of the top 30 proteins by fluorescent intensity from each study differ according to the normalization and pre-processing steps used and the ranking methods employed. Specifically, the displays compare the ranks of the top 30 proteins, based on fluorescent intensity, published in Kobayashi *and others* (2019) (Published Ranks) to those obtained by ranking point estimates of the posterior median fluorescent intensity of normalized signal from each protein across all arrays published in Bérubé *and others* (2021) (Pre-Processed ranks). We further compare these to the squared error loss ranks (Method A SEL) denoted as 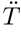 and the *γ*-ranks denoted as 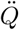 (Method A Gamma). In their study, Pan *and others* (2017) do not rank proteins by fluorescent intensity therefore we use the same procedures as for the malaria arrays and compare the squared error loss ranks and the *γ*-ranks from the HuProt^*T M*^ arrays to the Pre-Processed ranks based on point estimates. In both cases, however, the use of the full posterior distributions to produce optimal ranks under different loss functions substantially changes the ordering and composition of the top 30 proteins list. The statistical literature shows that ranking methods accounting for uncertainty by optimizing a loss function outperform other methods of ranking, (Shen and Louis, 1998; Lin *and others*, 2006; Henderson and Newton, 2016) therefore ranks for the malaria and lung cancer studies obtained by using the Bayseian model outputs are preferable to those obtained using other estimates of normalized fluorescent intensity (Bérubé *and others*, 2022).

**Fig. 6:**
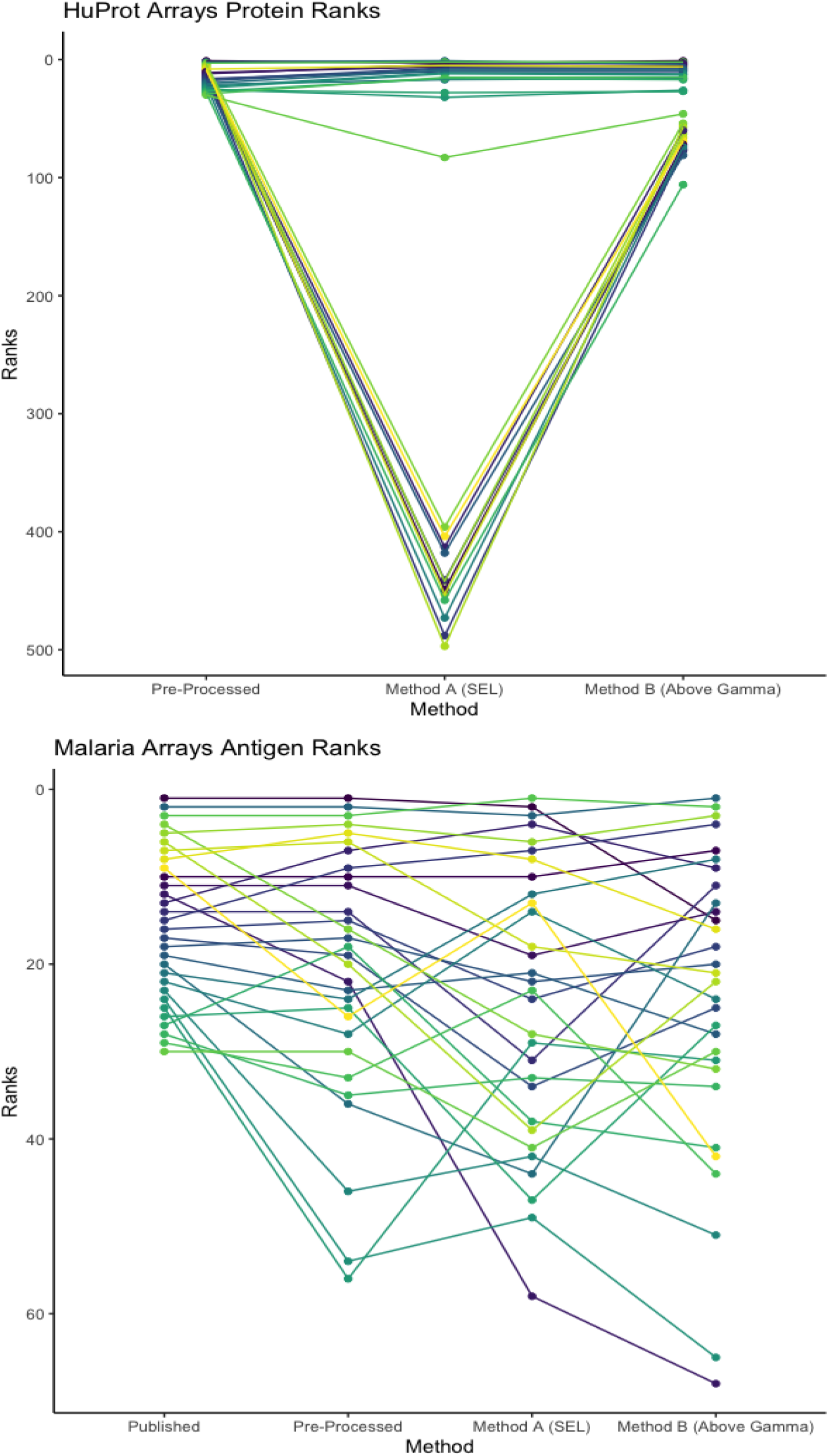
Each color is a protein on the original malaria or HuProt^*T M*^ array, each column shows one of: the published ranks from Kobayashi *and others* (2019), the ranks of fluorescent intensity pre-processed according to the procedure described in Bérubé *and others* (2021), the squared error loss ranks 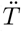, and the gamma ranks 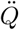. Lines represent how ranks of proteins change across each method of pre-processing and ranking.

## 4. Discussion

Protein microarray data have the ability to provide insight into important biological processes and to guide clinical and public health policy. However, high levels of between-array variation even after pre-processing complicate the comparison of protein levels across samples in a way that supports biological inference and quantification of uncertainty. For this reason, relative quantities of proteins, or array-specific ranks of proteins based on fluorescent intensity are necessary. As are all inferences, ranks are sensitive to pre-processing and normalization methods, but ranking approaches that accommodate uncertainty, especially in the Bayesian framework, produce loss-function, optimal ranks that can be used for a variety of downstream analyses.

We developed and applied a Bayesian model that extracts full posterior distributions of normalized fluorescent intensity for every protein in every serum sample of a protein microarray study. We showed that this model produces estimates, including ranks, that support downstream inferences. Simplifications to the full model substantially decreased computation time and fit equally well to the two datasets. We then evaluated the validity via a simulation study. We showed that the model has built-in flexibility that can accommodate substantially data different protein microarrays. The full posterior distributions obtained from the models can be used as inputs for variety of ranking methods that incorporate uncertainty (Laird and Louis, 1989; Shen and Louis, 1998; Lin *and others*, 2006; Henderson and Newton, 2016). Finally, we showed how the ranks of proteins and therefore subsequent analysis can be impacted by the choices of both pre-processing and ranking methods. Further investigation of the squared error loss ranks for antigens measured by the malaria arrays analyzed in Kobayashi *and others* (2019) revealed potential novel biomarkers of the timing and duration of past malaria exposure in adult populations.

In spite of the differences in manufacturing and features of the arrays used in the two datasets, our model fits quite well to both, suggesting the potential for it to be effective across multiple protein microarray studies. However, the two published studies represent a fairly narrow sample of the currently available protein microarrays, and the amount of between-array variability present is dependent at least in part on the manufacturing process. Therefore, a more comprehensive evaluation of our model on different types of protein microarrays is warranted.

Although investigation of percentile residuals for both datasets reveals a good fit for both the complete and simplified models, the data has larger spread than estimated by the prediction models, these discrepancies should be further examined but do not rise to the levels observed in the residual analysis of simulated data sets *C* and *F*. Therefore it is likely that the squared error loss ranks (SEL) of proteins in both datasets remain good estimates of true ranks. Generally, the influence of more extreme data values on model fit should be examined; possibly, some extreme values should be set aside during model fitting.

Appropriate pre-proccessing is an important component of any bioinformatic pipeline, however, after appropriate pre-processing, the use of estimates that propagate appropriate levels of uncertainty can support a wide range of inferential goals. Bayesian models enable analysts to accomplish these goals while providing other advantages, for instance, the use of full posterior distributions can be used to obtain optimal ranks. This is especially pertinent for protein microarrays, where ranks are the basis for most valid biological inference. While Bayesian models have been developed and evaluated for similar assays such as DNA microarrays, these models rely on assumptions that are inappropriate for protein microarray studies. Our model is developed specifically for protein microarrays and therefore has the potential to improve the robustness and accuracy of conclusions drawn from these data by propagating appropriate levels of uncertainty through bioinformatic pipelines.

## 5. Software

R code for all analyses and simulation can be found in the following GitHub repository: https://github.com/sberube3/Protein_array_bayesian_model.

## 6. Supplementary Material

Supplementary materials which include estimation methods for certain parameters, as well as plots that demonstrate the convergence and updating of various parameters and hyperparameters in the model are available online at http://biostatistics.oxfordjournals.org.

## Acknowledgments

The authors are grateful to Philip Felgner and D.Huw Davies for sharing data relating to their *Plasmodium falciparum* and *P. vivax* antibody arrays. The authors also acknowledge Jianbo Pan and Heng Zhu for sharing the protein microarray data from their lung cancer study. The authors gratefully acknowledge partial support from the following sources: NIH-NIAID, U19-AI089680.

