## Supplement for "A Bayesian Hierarchical Model for Signal Extraction from Protein Microarrays"

February 16, 2022

#### 1 Appendix A: Array Composition

**Table S 1:** Values of $t(p)$ on the HuProt$^{TM}$ array.

| Protein Category | Protein ($p(i,j)$) | Number of Spots ($j$) | Type t(p) |
|---|---|---|---|
| Negative Control | GST 10 ng/$\mu$l | 48 | 1 |
| | GST 50 ng/ $\mu$l | 48 | 2 |
| | GST 100 ng/$\mu$l | 48 | 3 |
| | GST 200 ng/$\mu$l | 48 | 4 |
| | Mouse-anti-biotin | 48 | 5 |
| | Rabbit-anti-biotin | 48 | 6 |
| | BSA | 48 | 7 |
| | Buffer | 816 | 8 |
| | Empty | 3072 | 9 |
| Positive Control | Histone 1 | 48 | 10 |
| | Histone 2 (A+B) | 48 | 11 |
| | Histone 3 | 48 | 12 |
| | Histone 4 | 48 | 13 |
| | Alexa Fluor labeled IgG | 48 | 14 |
| | Rhodamine+ Alexa Fluor labeled IgG | 48 | 15 |
| | Biotin-BSA | 48 | 16 |
| | Mouse IgM | 48 | 17 |
| Active Proteins | Active Proteins | 48384 | $a_1$ |

**Table S 2:** Values of $t(p)$ on the malaria array.

| Control Type | Protein $p(i,j)$ | Number of Spots $j$ | Type t(p) |
|---|---|---|---|
| Negative Control | Blank1 | 4 | 1 |
| | Blank2 | 5 | 2 |
| | Empty | 4 | 3 |
| | No DNA Reaction | 24 | 4 |
| | TTBS | 32 | 5 |
| Positive Control | anti-human IgG 0.003 | 4 | 6 |
| | anti-human IgG 0.03 | 4 | 7 |
| | anti-human IgG 0.3 | 4 | 8 |
| | anti-human IgG 0.001 | 4 | 9 |
| | anti-human IgG 0.01 | 4 | 10 |
| | anti-human IgG 0.1 | 4 | 11 |
| | IgG mix 0.003 | 4 | 12 |
| | IgG mix 0.03 | 4 | 13 |
| | IgG mix 0.3 | 4 | 14 |
| | IgG mix 0.001 | 4 | 15 |
| | IgG mix 0.01 | 4 | 16 |
| | IgG mix 0.1 | 4 | 17 |
| Active Proteins | Active Proteins | 1038 | $a_1$ |

### 2 Appendix B: Estimating the error distribution

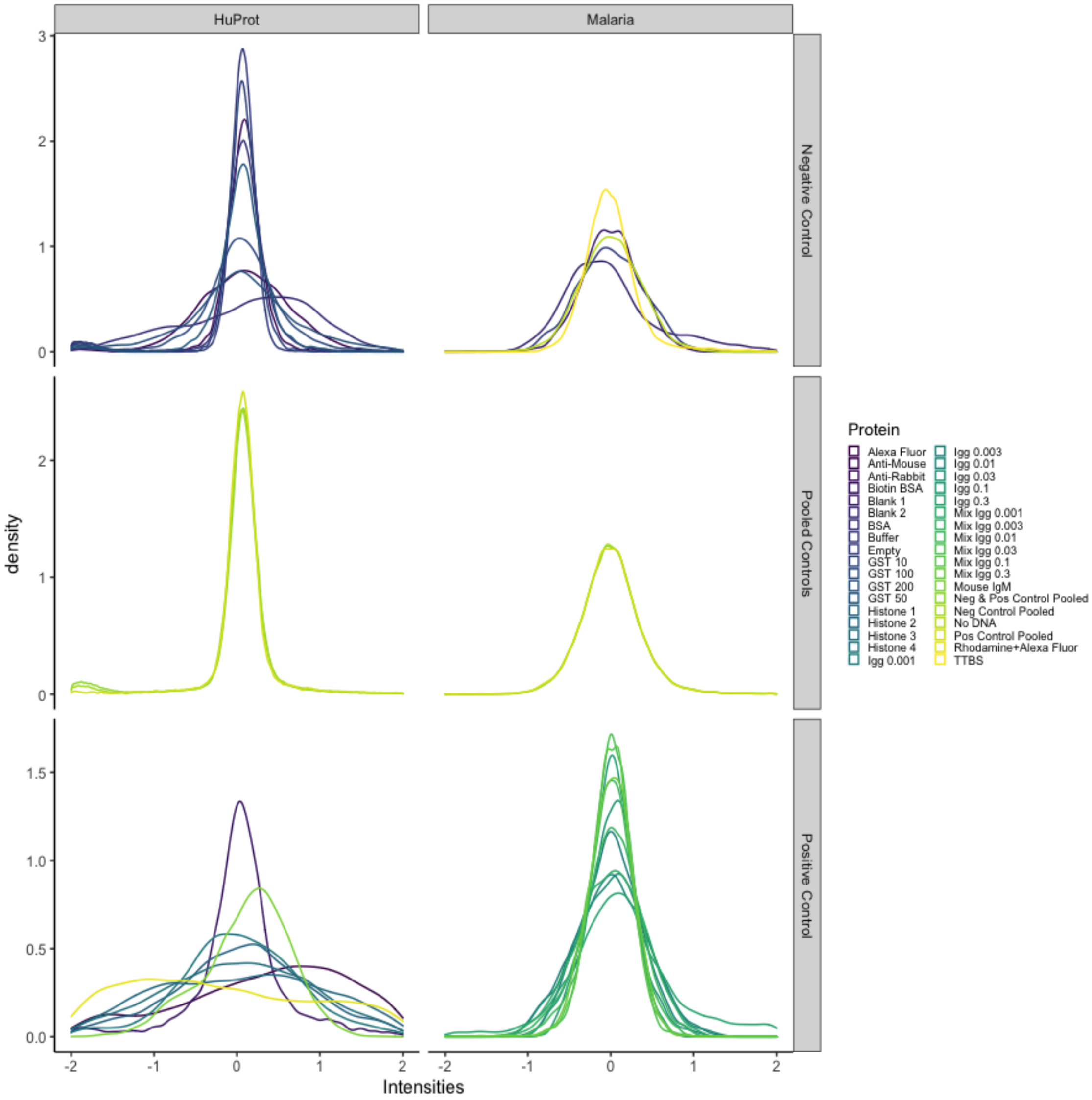


**Fig S 1:** Observed distributions of $e_{i,t(p),p(i,j),j}$ for negative, positive, and combined positive and negative controls.

#### 2.1 Estimating $\alpha$

To estimate $\alpha$, we will use a maximum likelihood estimation procedure that uses an empirical estimate of the error distributions $e_{i,t(p),p(i,j),j}$ for $t(p) \in \{1, ..., T\}$. According to our proposed model in Equation 2.1 (main text), the distribution of $e_{i,t(p),p(i,j),j}$ across all arrays can be estimated from the data by considering the distribution standardized control probes $Y_{i,t(p),p(i,j),j} - \hat{\mu}_{i,t(p),p(i,j)}$, where $\hat{\mu}_{i,t(p),p(i,j)}$ is the sample mean taken over all spots of type $t(p), \in \{1, ..., T\}$ on each array. We obtain an approximate maximum likelihood estimate (MLE) for $\alpha$ based on the following likelihood:

$$L(Y_{i,t(p),p(i,j),j}|\alpha) = \prod_j \frac{1}{B(\alpha,\alpha)} \times \Phi\left(\frac{y_{i,t(p),p(i,j),j} - \hat{\mu}_{i,t(p),p(i,j)}}{\hat{\sigma}} c(\alpha)\right)^{\alpha-1} \times \left[1 - \Phi\left(\frac{y_{i,t(p),p(i,j),j} - \hat{\mu}_{i,t(p),p(i,j)}}{\hat{\sigma}} c(\alpha)\right)\right]^{\alpha-1} \times \phi\left(\frac{y_{i,t(p),p(i,j),j} - \hat{\mu}_{i,t(p),p(i,j)}}{\hat{\sigma}} c(\alpha)\right) \times \frac{c(\alpha)}{\hat{\sigma}} \quad (1)$$

Where $\hat{\sigma}$ is the sample standard deviation of the set $Y_{NC}$, or all observations of negative control probes across all arrays. Given that it is not possible to obtain a closed form equation for the maximum likelihood estimate (MLE) of $\alpha$ with the likelihood in Equation 1, we propose a numerical estimation that uses a fine-grained "grid" of possible $\alpha$ values ranging from $1 - 17$ and increasing by increments of $\frac{1}{1000}$ to identify the value of $\alpha$ that produces the highest likelihood value. We replicate this estimation procedure for positive control proteins and then for the combined positive and negative control proteins. Figure 2 shows the log likelihood values of equation 1 for each of the three sets of proteins in the malaria and Huprot$^{TM}$ arrays. The value of $\alpha$ that produces the maximum log likelihood value is shown with a red vertical line.

### 3 Appendix C: Posterior Distributions and Trace Plots

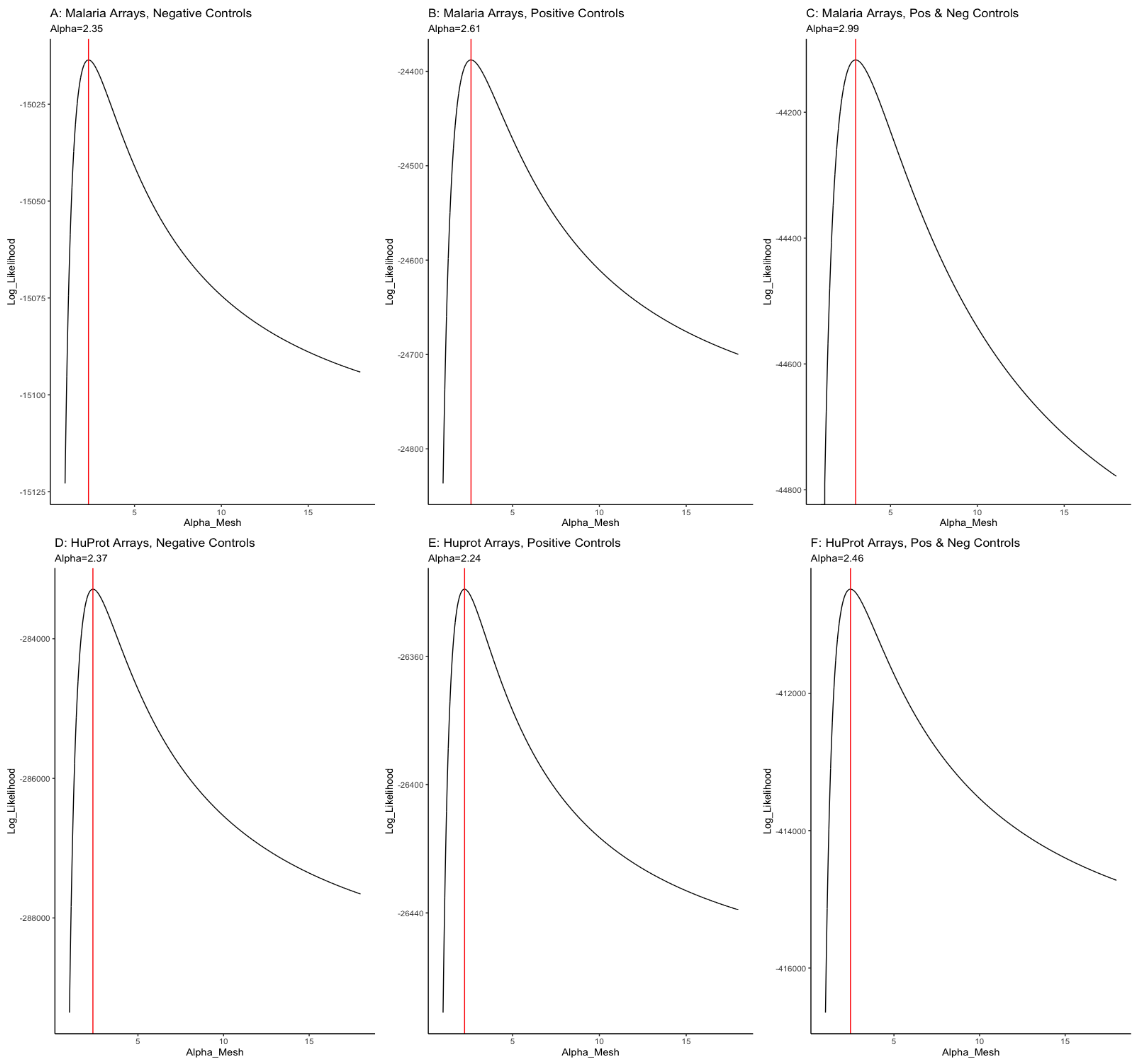


**Fig S 2:** Log likelihood values of equation 1 for each of the three sets of proteins in the malaria arrays, negative control (A and D), positive control (B and E) and combined positive and negative control (C and F) proteins across values of $\alpha$ in the grid. The value of $\alpha$ that produces the maximum log likelihood value is shown with a red vertical line.

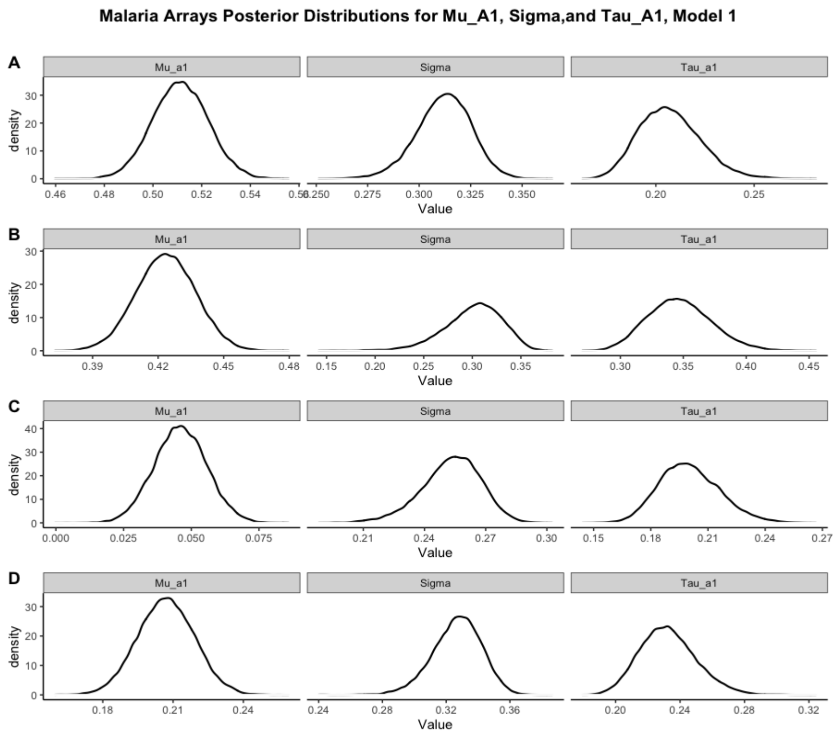


**Fig S 3:** Posterior distributions for $\sigma_i$, $\mu_{i,a_1}$, and $\tau_{i,a_1}$ for four malaria (A,B,C,D) arrays fit with Model 1.

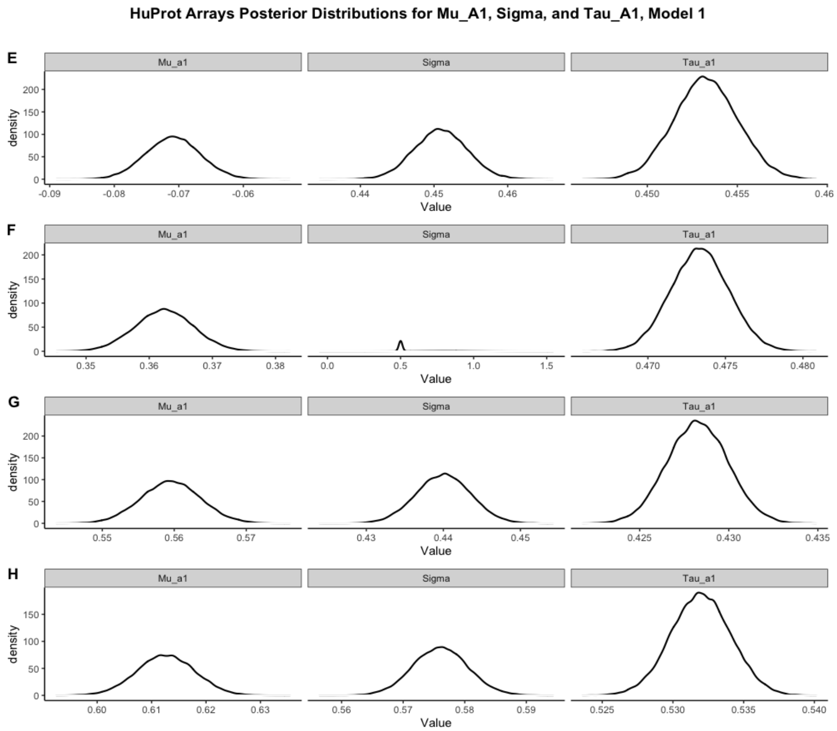


**Fig S 4:** Posterior distributions for $\sigma_i$, $\mu_{i,a_1}$, and $\tau_{i,a_1}$ for four HuProt$^{TM}$ (E,F,G,H) arrays fit with Model 1.

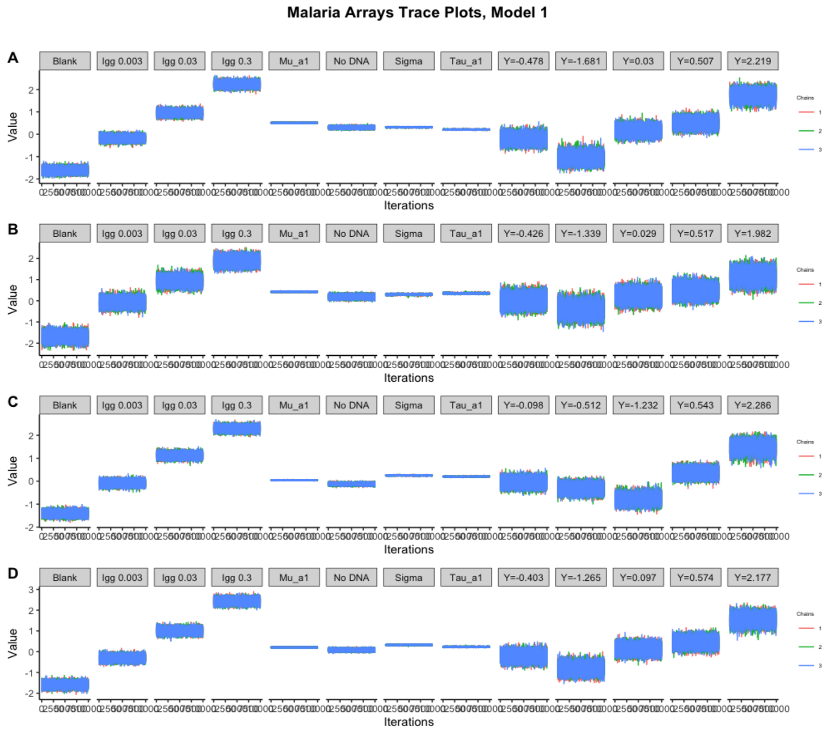


**Fig S 5:** Trace plots for estimated parameters, $S, \mu_{i,t(p)}, \mu_{i,a_1}, \tau_{i,a_1}, \sigma_i$ obtained by fitting Model 1 to four malaria arrays (A,B,C,D).

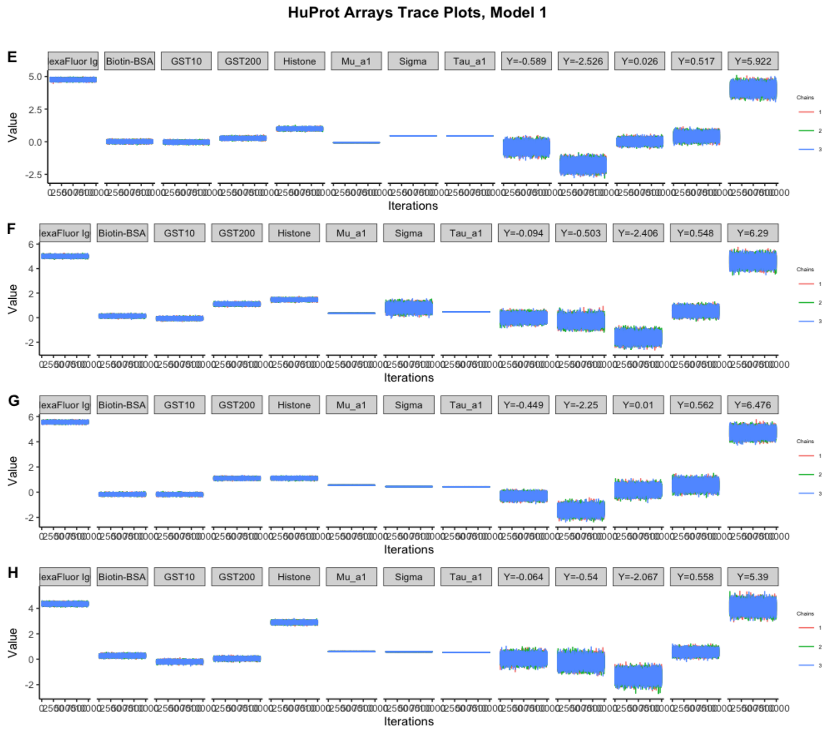


**Fig S 6:** Trace plots for estimated parameters, $S, \mu_{i,t(p)}, \mu_{i,a_1}, \tau_{i,a_1}, \sigma_i$ obtained by fitting Model 1 to four HuProt$^{TM}$ (E,F,G,H) arrays.

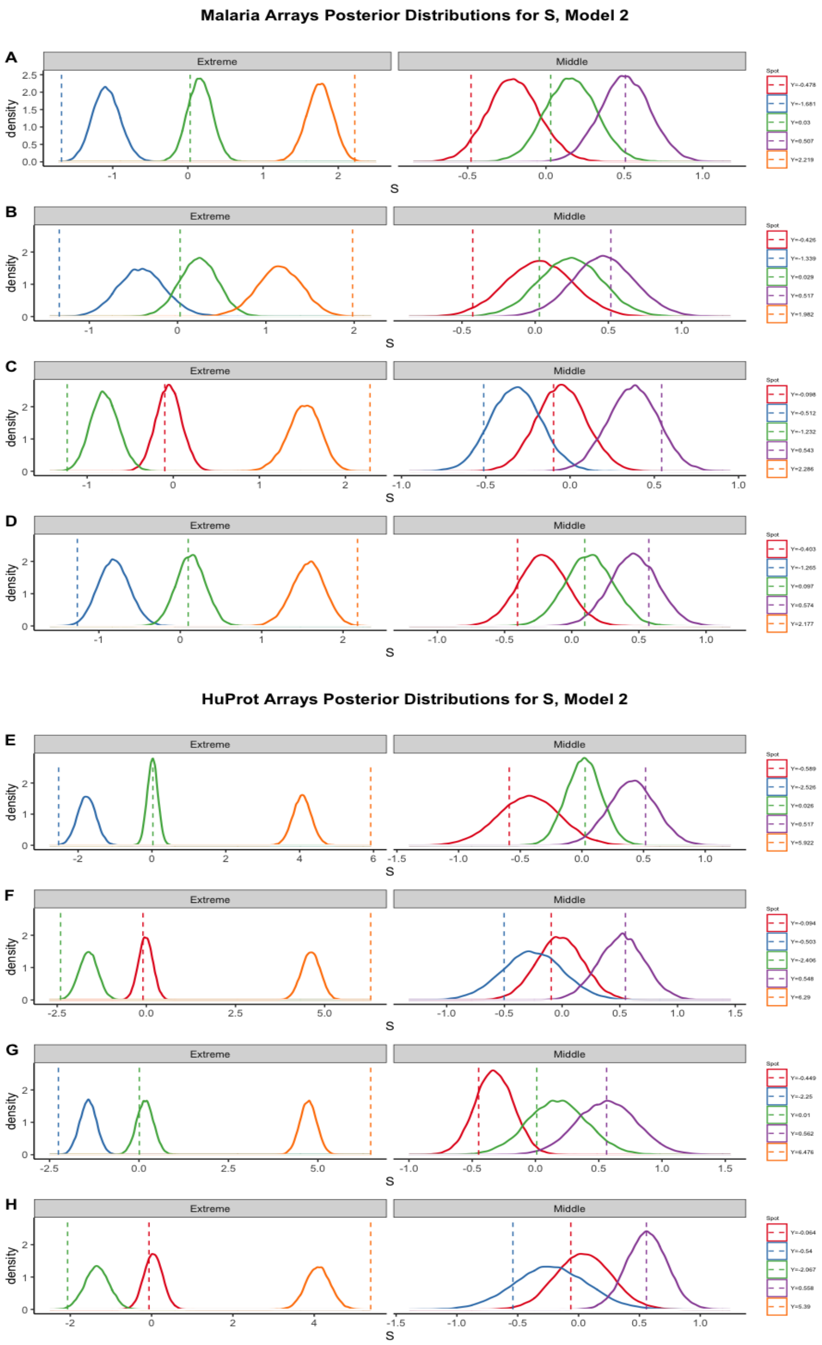


**Fig S 7:** Posterior distributions for $S_{i,t(p),p(i,j),j}$ for five different proteins on four malaria (A,B,C,D) and HuProt$^{TM}$ (E,F,G,H) arrays fit with Model 2. Vertical dashed lines are at the value or means of values $Y_{i,t(p),p(i,j),j}$ corresponding to the protein $p(i,j)$.

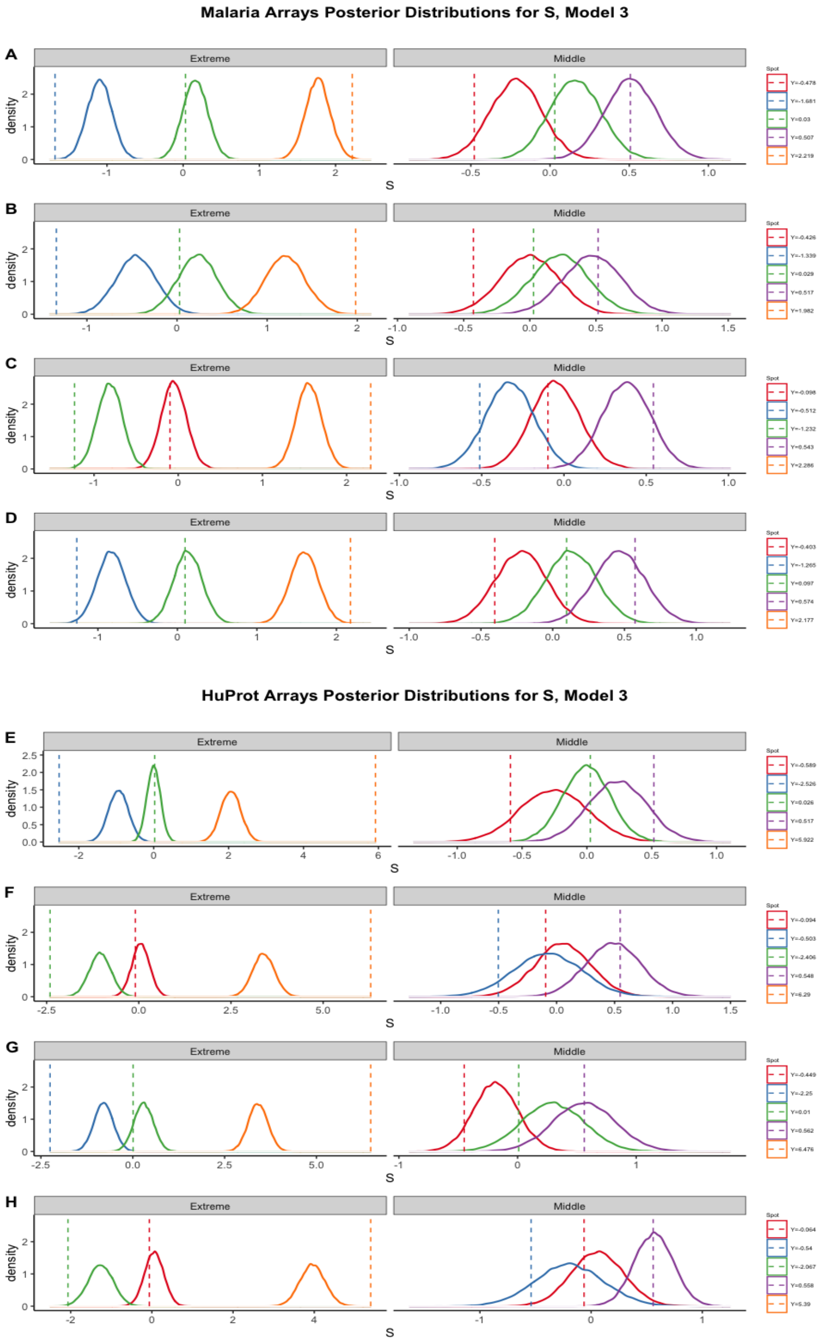


**Fig S 8:** Posterior distributions for $S_{i,t(p),p(i,j),j}$ for five different proteins on four malaria (A,B,C,D) and HuProt$^{TM}$ (E,F,G,H) arrays fit with Model 2. Vertical dashed lines are at the value or means of values $Y_{i,t(p),p(i,j),j}$ corresponding to the protein $p(i,j)$.

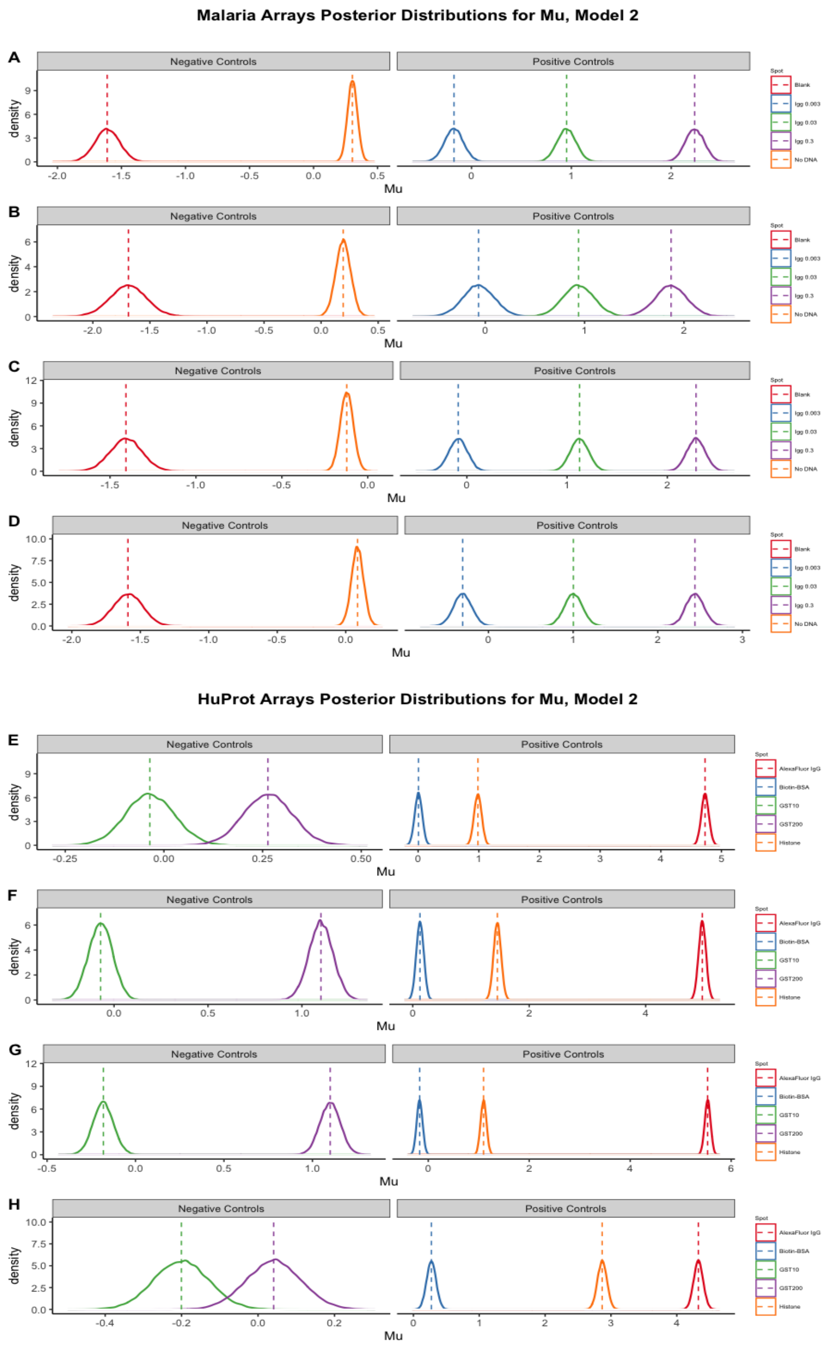


**Fig S 9:** Posterior distributions for $\mu_{i,t(p)}$ for five different types $t(p)$ if control proteins on four malaria (A,B,C,D) and HuProt$^{TM}$ (E,F,G,H) arrays fit with Model 2. Vertical dashed lines are at the means of values $Y_{i,t(p)j}$ corresponding to the proteins with type $t(p)$.

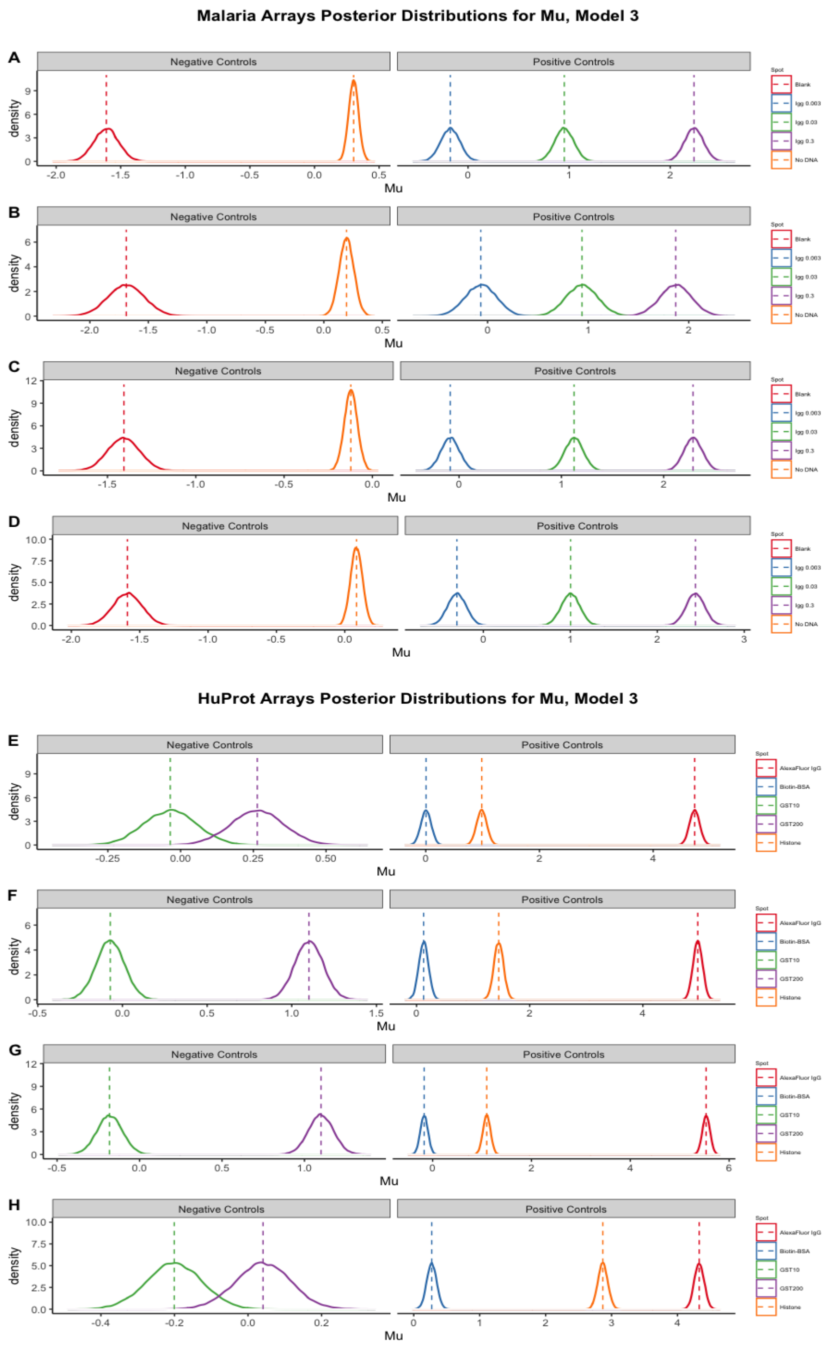


**Fig S 10:** Posterior distributions for $\mu_{i,t(p)}$ for five different types $t(p)$ if control proteins on four malaria (A,B,C,D) and HuProt$^{TM}$ (E,F,G,H) arrays fit with Model 3. Vertical dashed lines are at the means of values $Y_{i,t(p)j}$ corresponding to the proteins with type $t(p)$.

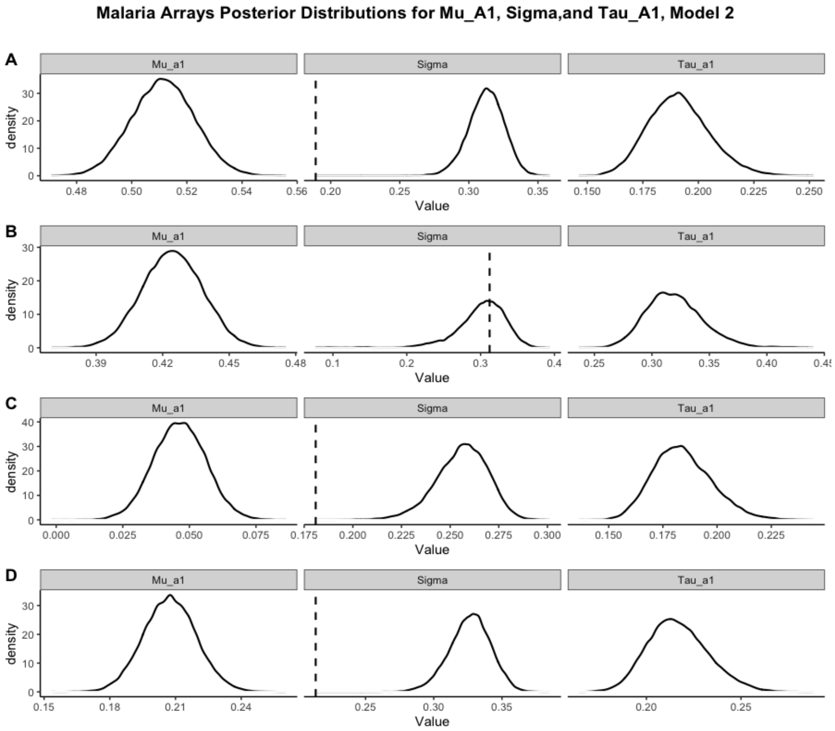


**Fig S 11:** Posterior distributions for $\sigma_i$, $\mu_{i,a_1}$, and $\tau_{i,a_1}$ for four malaria (A,B,C,D) arrays fit with Model 2. Vertical dashed lines are at the value $\hat{\sigma}_i$.

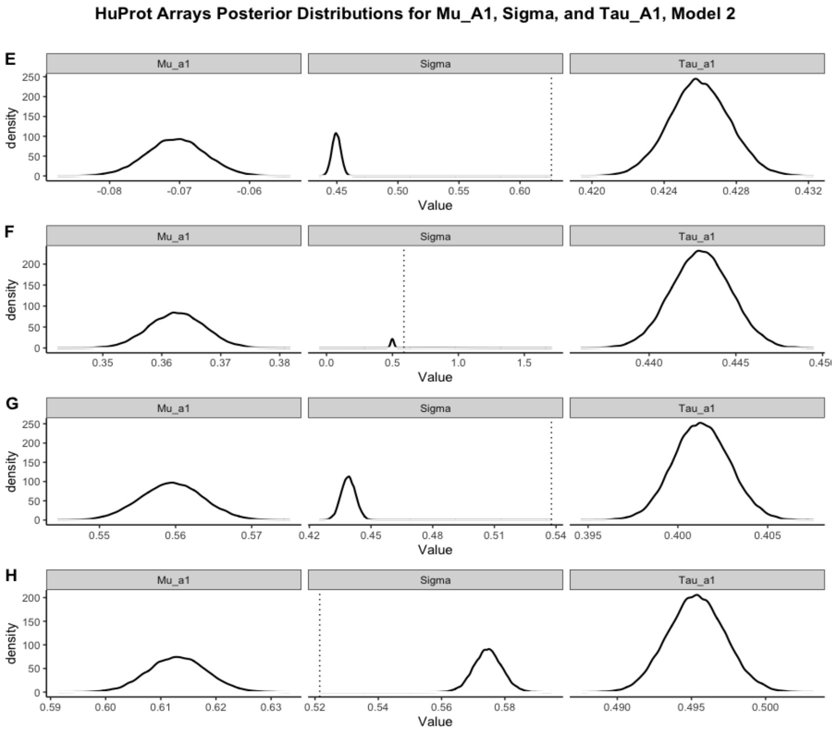


**Fig S 12:** Posterior distributions for $\sigma_i$, $\mu_{i,a_1}$, and $\tau_{i,a_1}$ for four HuProt$^{TM}$ (E,F,G,H) arrays fit with Model 2. Vertical dashed lines are at the value $\hat{\sigma}_i$.

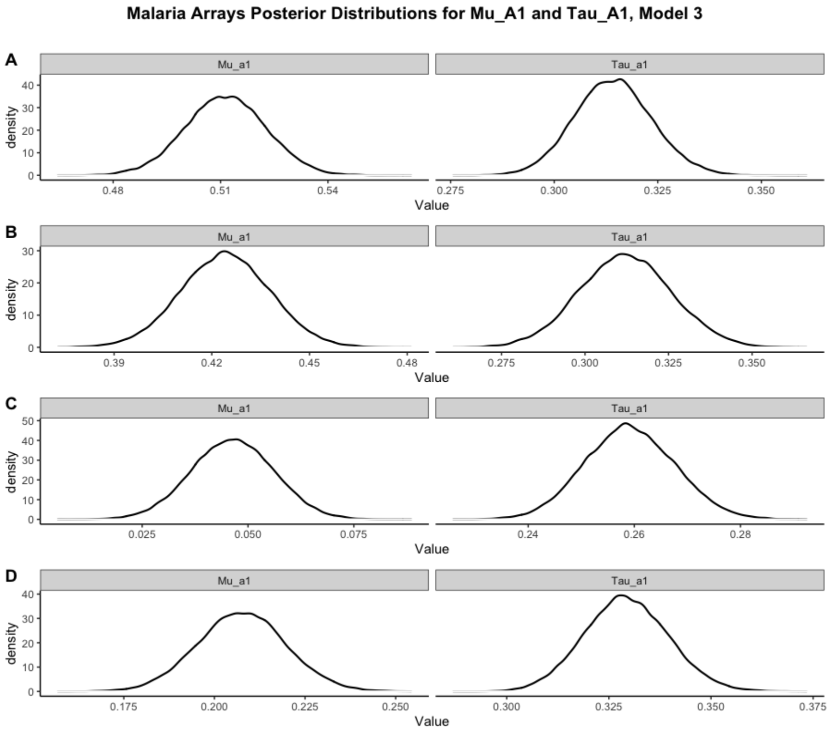


**Fig S 13:** Posterior distributions for $\mu_{i,a_1}$, and $\tau_{i,a_1}$ for four malaria (A,B,C,D) arrays fit with Model 3.

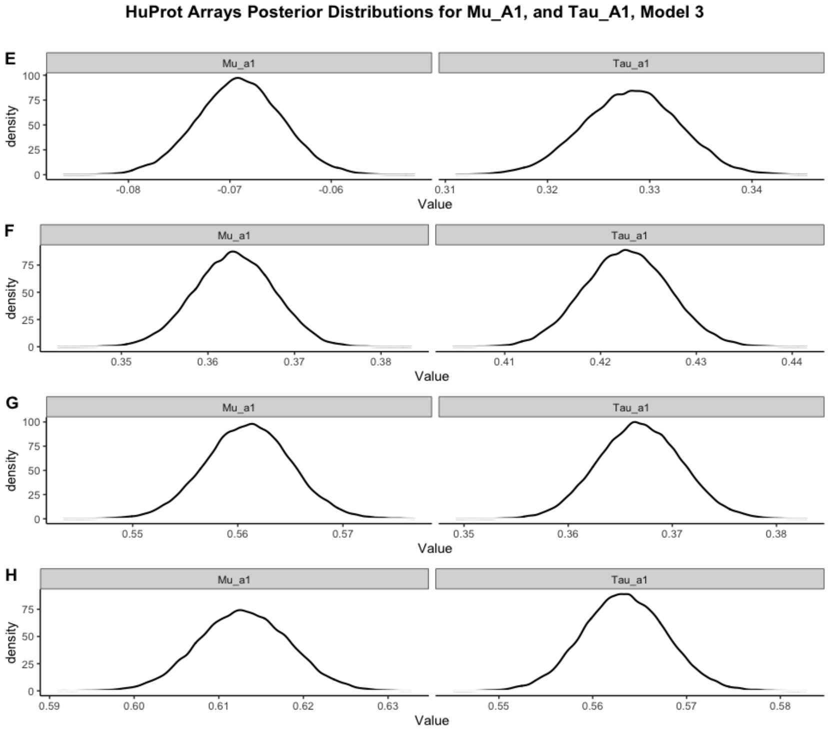


**Fig S 14:** Posterior distributions for $\mu_{i,a_1}$, and $\tau_{i,a_1}$ for four HuProt$^{TM}$ (E,F,G,H) arrays fit with Model 3.

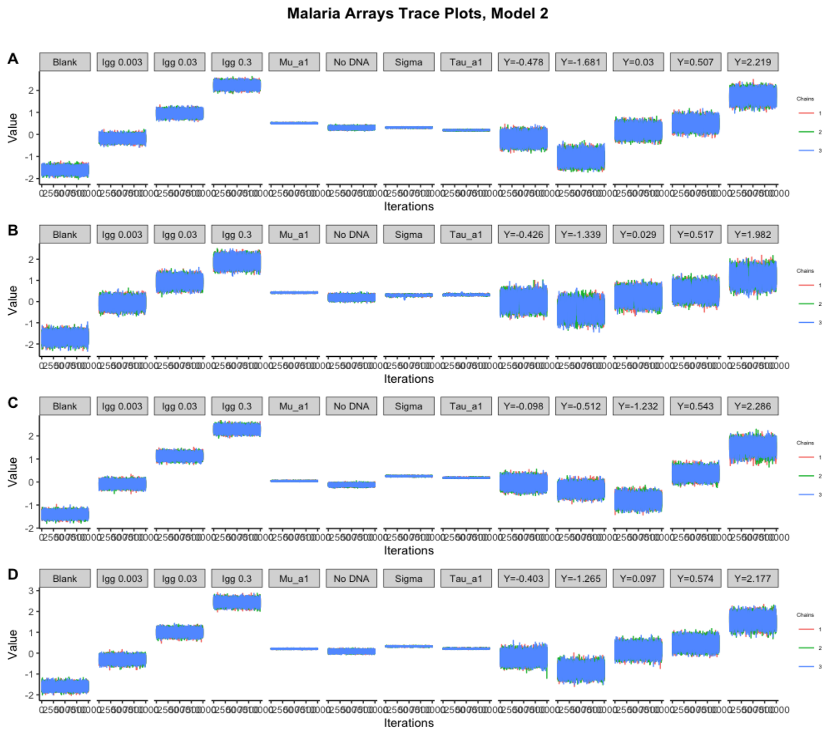


**Fig S 15:** Trace plots for estimated parameters, $S, \mu_{i,t(p)}, \mu_{i,a_1}, \tau_{i,a_1}, \sigma_i$ obtained by fitting Model 2 to four malaria arrays (A,B,C,D).

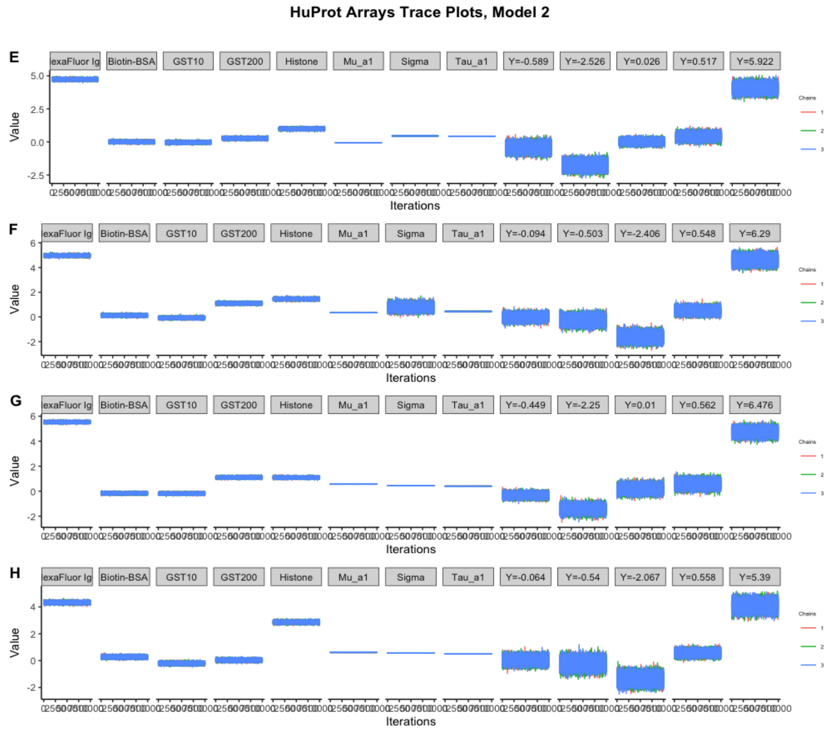


**Fig S 16:** Trace plots for estimated parameters, $S, \mu_{i,t(p)}, \mu_{i,a_1}, \tau_{i,a_1}, \sigma_i$ obtained by fitting Model 2 to four HuProt$^{TM}$ (E,F,G,H) arrays.

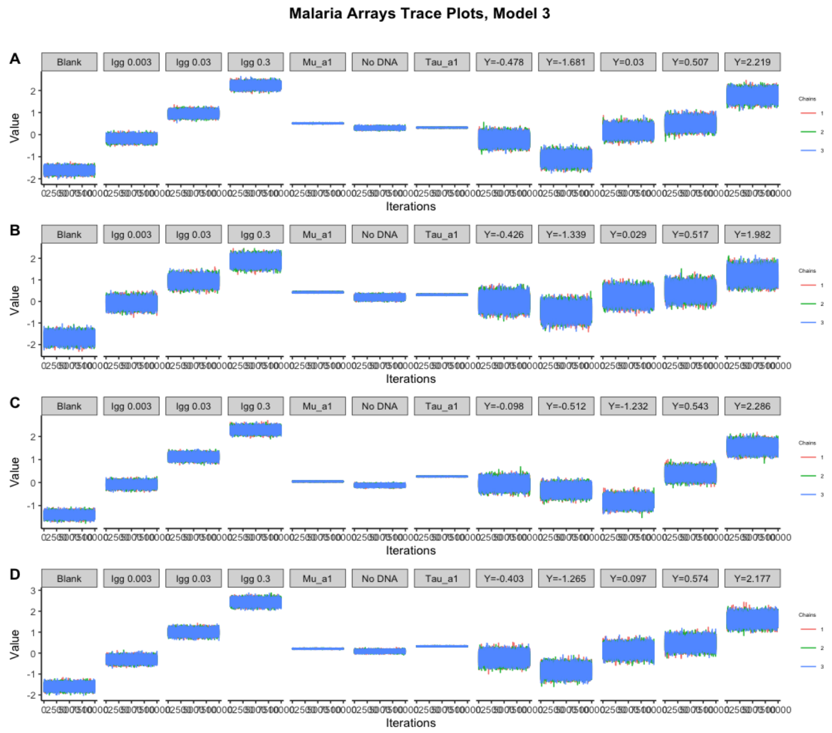


**Fig S 17:** Trace plots for estimated parameters, $S, \mu_{i,t(p)}, \mu_{i,a_1}, \tau_{i,a_1}, \sigma_i$ obtained by fitting Model 3 to four malaria arrays (A,B,C,D).

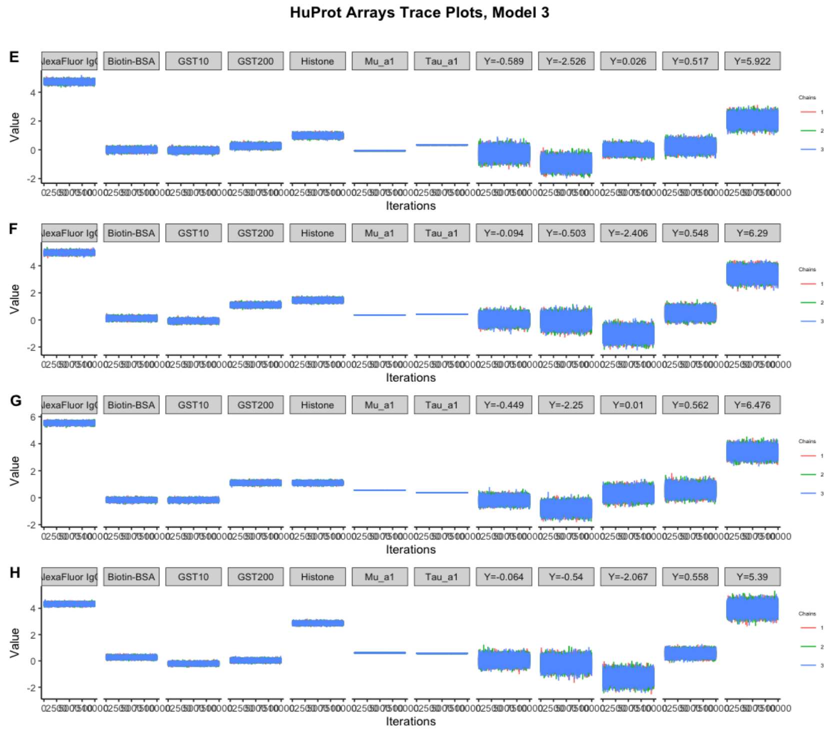


**Fig S 18:** Trace plots for estimated parameters, $S, \mu_{i,t(p)}, \mu_{i,a_1}, \tau_{i,a_1}, \sigma_i$ obtained by fitting Model 3 to four HuProt$^{TM}$(E,F,G,H) arrays.
